# Iron Metabolism as a Therapeutic Vulnerability in Stem Cell-Like Castration-Resistant Prostate Cancer

**DOI:** 10.64898/2026.02.24.707686

**Authors:** Wanli Cheng, Andrea Brunello, Francesco Bonollo, Thomas Michael Marti, Panagiotis Chouvardas, David P Labbé, Marta De Menna, George Thalmann, Sofia Karkampouna, Marianna Kruithof-de Julio

## Abstract

Prostate cancer is the second most common malignancy among men, with androgen deprivation therapy (ADT) serving as the standard treatment due to the hormone sensitivity of prostate tumors. However, therapeutic resistance frequently develops, leading to castration-resistant prostate cancer (CRPC), an aggressive and lethal disease. A recently defined subtype, stem cell-like CRPC (CRPC-SCL), accounts for approximately 25% of CRPC cases and demonstrates poor responsiveness to ADT. CRPC-SCL is characterized by the expression of Cluster of Differentiation 44 (CD44), a glycoprotein that promotes hyaluronic acid binding and uptake. Within CRPC-SCL patient-derived xenograft (PDX) model, CD44 high (CD44hi) cells exhibit enhanced tumorigenicity and proliferative capacity. Importantly, iron metabolism emerges as a critical regulator of this population: CD44hi cells maintain elevated intracellular iron, which sustains CD44 expression and stem cell-like properties by modulating H3K9me2 modification. Leveraging this vulnerability, inhibition of the iron-regulatory factor NRF2 was shown to increase intracellular free iron and selectively induce ferroptosis in CD44hi cells. These findings highlight the therapeutic potential of targeting iron metabolism to induce ferroptosis as a novel treatment strategy for CRPC-SCL.

## Introduction

Prostate cancer (PCa) is the second most common cancer among men worldwide [1]. Androgen deprivation therapy (ADT) is commonly used to inhibit tumor growth due to the hormone sensitivity of the prostate [2]. However, a subset of tumors eventually develops resistance and progresses to castration-resistant prostate cancer (CRPC) [3]. Recent research has classified a novel stem cell-like subtypes (SCL) of CRPC based on chromatin profiling [4]. The CRPC-SCL subtype, which accounts for approximately 30% of CRPC cases, is particularly challenging due to its poor response to conventional ADT and its association with aggressive clinical behavior [4]. This finding highlights the pressing need to develop subtype-specific therapeutic strategies for CRPC-SCL.

CD44, a cell-surface glycoprotein, mediates the binding and uptake of hyaluronic acid (HA) [5]. The CD44 is highly and specifically expressed in the CRPC-SCL subtype, suggesting its potential as a marker for this lineage [4]. Previous studies have identified CD44 as a cancer stem cell marker in primary PCa, capable of surviving castration and contributing to the development of CRPC [6]. These observations indicate that investigating the role of CD44 may provide important insights into the biology of CRPC-SCL and offer new avenues for therapeutic intervention.

Previous studies have reported that iron and copper can bind to HA to facilitate uptake via CD44 [7]. Iron is an essential biomineral with critical roles in electron transport, oxygen delivery, and the maintenance of redox homeostasis in mammalian cells [8]. Beyond these canonical functions, iron has been shown to promote demethylation through activation of the Jumonji C (JMJC) family of enzymes [9]. Importantly, iron also underlies a distinct form of regulated cell death known as ferroptosis, which is characterized by the accumulation of lipid peroxides and oxidative stress driven by iron overload [10]. Ferroptosis can be initiated by antioxidant depletion, iron dysregulation, and alterations in lipid metabolism, and accumulating evidence suggests that it may represent a promising therapeutic strategy for prostate cancer [11]. However, the role of CD44 in CRPC-SCL remains unclear, and it is not yet known whether iron metabolism contributes to CD44 function and CRPC-SCL biology. Addressing this question may not only improve our understanding of CRPC-SCL but also inform the development of subtype-specific therapeutic approaches.

In this study, we demonstrate that CD44hi cells exhibit pronounced SCL features and enhanced tumorigenic potential in CRPC. Furthermore, we identify iron as a key regulator of CD44 expression and SCL characteristics through epigenetic mechanisms. Importantly, targeting iron metabolism by inhibiting NRF2 to induces ferroptosis in CD44hi cells, thereby unveiling a potential therapeutic strategy for CRPC-SCL.

## Result

### CD44 is a putative marker of the CRPC-SCL subtype and represents a specific subpopulation within CRPC

Previous work has stratified CRPC into four distinct molecular subtypes [4]. Among these, the CRPC-SCL subtype displayed the highest iron score, reflecting heightened iron metabolic activity (Figure 1A), and was characterized by elevated expression of CD44, which has been proposed as a defining marker of this subtype (Figure 1B). To identify a representative model for the CRPC-SCL subtype, we performed RNA-seq profiling across our CRPC organoid models. Comparative analysis revealed that LAPC9 organoids exhibited transcriptomic features consistent with the SCL subtype (Figure 1C), whereas MSKPCa16 (PCa16) and WCM154 (PM154) aligned with Wnt and NE subtypes, respectively (Supplementary Fig. 1A-B), consistent with prior report [4]. Notably, AR activity scores in the LAPC9 model did not differ significantly from those in the NE model PM154 (Supplementary Fig. 1C), indicating that LAPC9 maintains low AR activity. In line with the SCL signature, LAPC9 displayed elevated CD44 expression together with enhanced iron metabolic activity, establishing it as a suitable model for studying the SCL subtype (Figure 1D-F). We next investigated the clinical relevance of CD44 expression in PCa. In the EMPACT cohort, immunohistochemical (IHC) analysis of tissue microarrays from primary PCa patients after radical prostatectomy, demonstrated that CD44 expression was associated with metastatic progression, supporting its role as an oncogenic driver (Figure 1G-H). To further delineate CD44-associated features, we interrogated single-cell RNA-seq data from CRPC patients [12, 13]. CD44 expression was enriched within a distinct cellular cluster of tumor cells from a CRPC patient, with CD44high (CD44hi) cells exhibiting stem cell-like properties (Supplementary Fig. 1E-H). Consistent findings were observed in an independent CRPC patient sample (Supplementary Fig. 1I-K), indicating that CD44 delineates a specific subpopulation within CRPC tumors. Together, these findings highlight LAPC9 as a model for the CRPC-SCL subtype and implicate CD44 as a potential driver of its aggressive biology.

**Fig 1.**
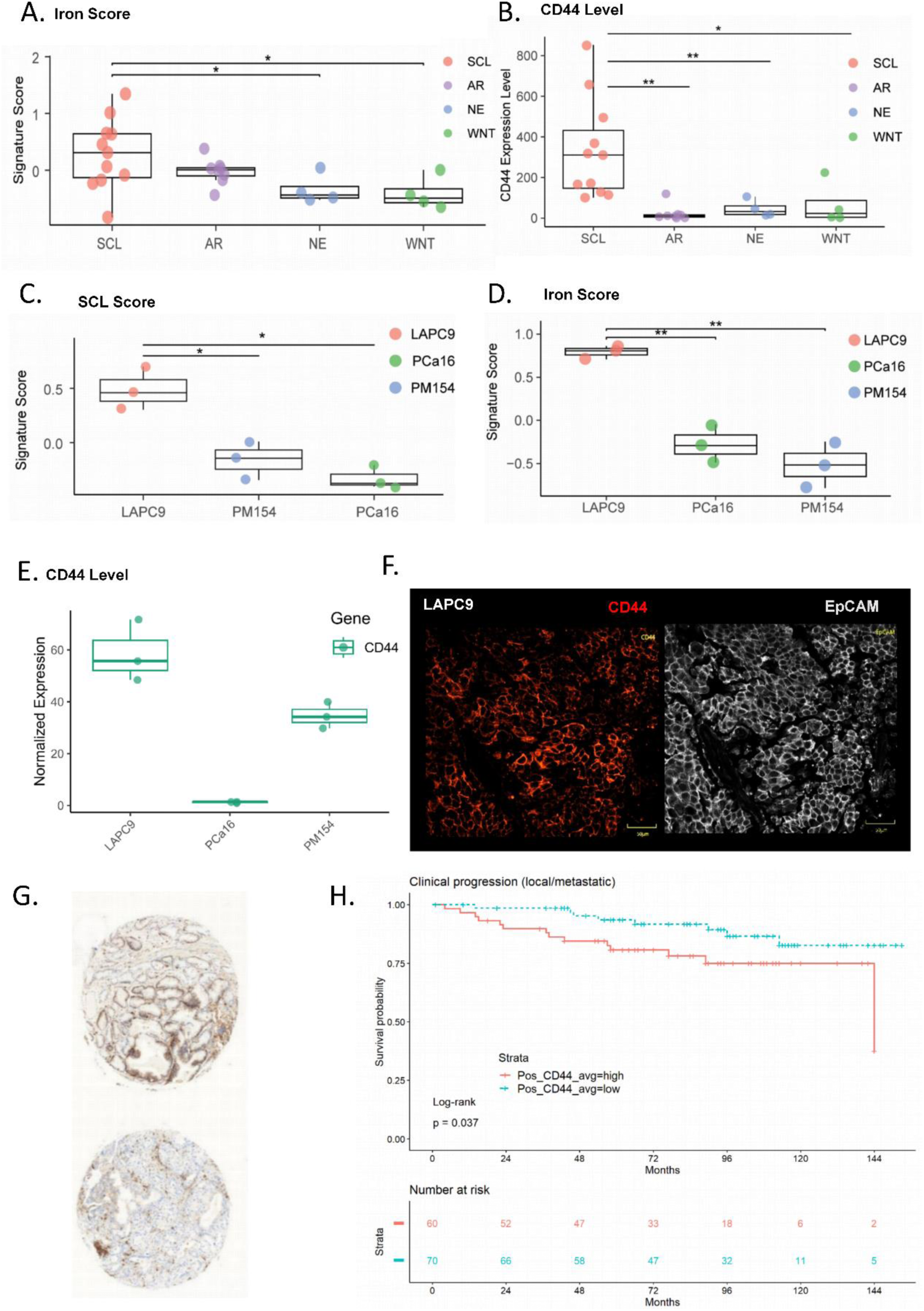
CD44 as distinct subpopulation in PCa. **A, B.** Iron signatures and CD44 level in different CRPC subtypes (androgen receptor (AR)-driven, Wnt-driven, neuroendocrine (NE), and stem cell-like (SCL) subtypes). **C, D.** Signature analysis identified SCL and iron scores base on RNA-seq data among LAPC9, PCa16 and PM154 organoids. **E.** CD44 expression among LAPC9, PCa16 and PM154 organoids. **F.** CD44 expression in LAPC9 by IF staining. **G-H.** The number of CD44-positive cells (protein expression) was evaluated in a tissue microarray (TMA) derived from primary prostate cancer patient samples in the EMPACT cohort by immunohistochemistry (IHC) in (**G**). The Kaplan–Meier (K-M) analyses of clinical progression based on expression levels of CD44 in TMA in(**H**). K-M curve compared by Log-rank test, p<0.05.

### The CD44hi subpopulation exhibited high tumorigenic potential both *in vitro* and *in vivo* in the CRPC-SCL model

In order to understand the role of CD44 subpopulation in SCL tumor, we separate the CD44hi and CD44low subpopulation by FACS sorting in LAPC9-copGFP-CBR tumor (Figure 2A). These sorted cells were maintained as organoid cultures as previously described by us [14]. The CD44hi subpopulation formed higher number and size organoids after 4 days culture compared to CD44low cells (Figures 2B-C). This demonstrated that CD44hi subpopulation have higher organoid-forming capacity compared to the CD44low subpopulation. To trace the dynamics of CD44hi and CD44low subpopulations *in vivo*, LAPC9 tumors were labeled with copGFP-CBR reporters, respectively. CD44hi and CD44low cells derived from LAPC9-copGFP tumor were injected into mice, and tumor burden was monitored weekly using IVIS-CT imaging (Figure 2D). Consistent with the *in vitro* findings, CD44hi cells exhibited enhanced tumor initiation capacity and significantly higher proliferation, resulting in larger tumors at the experimental endpoint (Figures 2E-F). Notably, tumors originating from CD44low cells transitioned to a CD44hi state to sustain tumor growth, whereas CD44hi-derived tumors consistently maintained high CD44 expression levels (Figure 2G-H). Together, these results establish CD44hi cells as a stable, highly tumorigenic subpopulation that drives organoid formation and tumor outgrowth, while CD44low cells display plasticity by converting to the CD44hi state *in vivo*.

**Fig 2.**
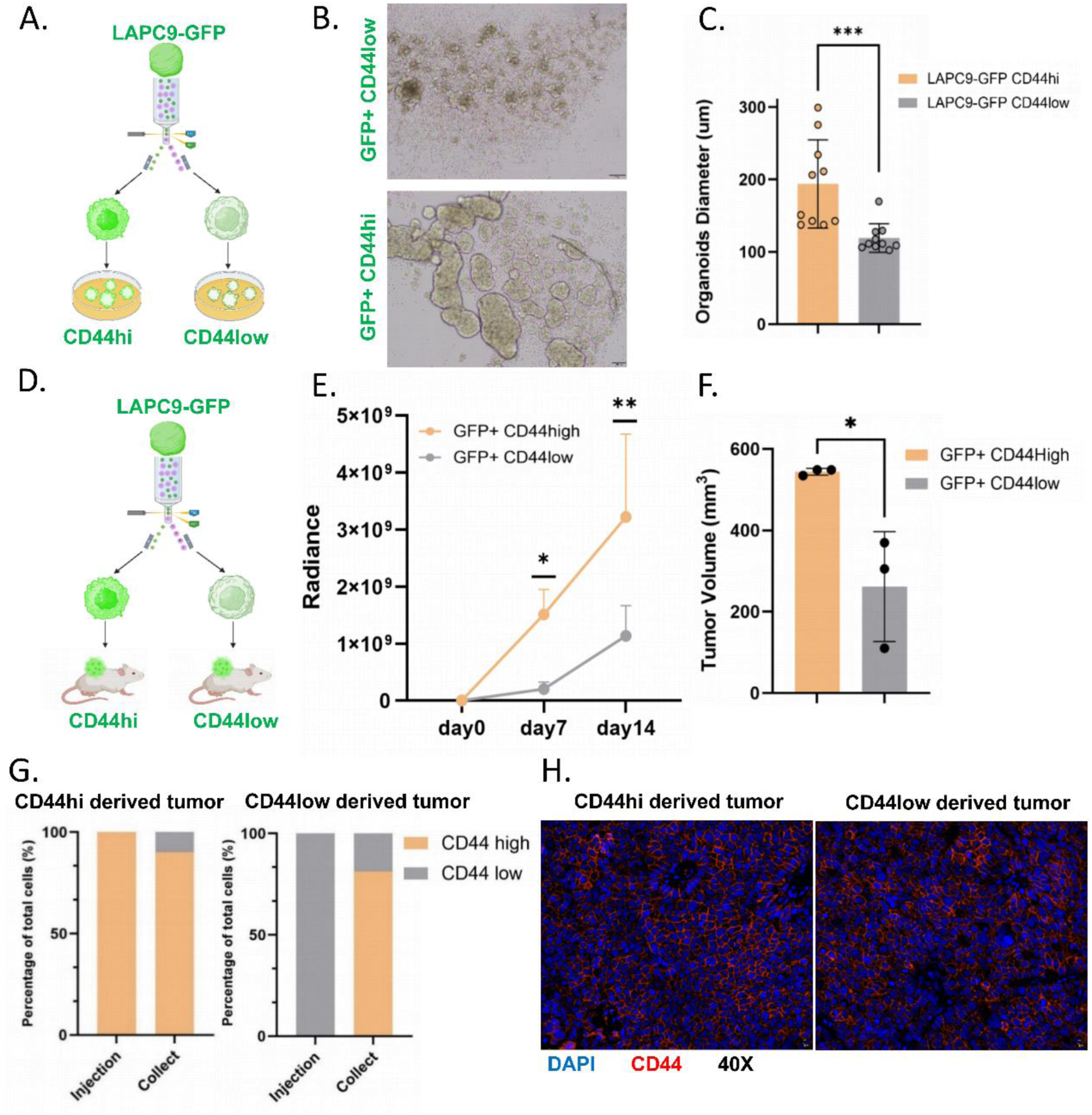
CD44hi subpopulation exhibited highly tumorigenicity. **A-C.** Fluorescence-activated cell sorting preformed on LAPC9-copGFP-CBR tumor to sort CD44hi and CD44low subpopulations (with GFP+ and luciferase). The sorted CD44hi cells and CD44low cells were cultured as organoids respectively. Size of organoids were quantified by W8 (n=10). **D.** The sorted CD44hi and low (with GFP+ and luciferase) cells were separately injected into mice (n=3 mice/group). **E.** Luciferase was measured by IVIS-CT in both CD44hi and CD44low groups weekly. The quantified radiance was used to measure tumor dynamics between CD44hi and low groups (n=3 mice). **F.** The tumors were collect at endpoint (day 46), tumor sizes were measured by scale (Hight*Length*Width*0.8), unpaired t-test for CD44hi and low groups (n=3 mice). **G.** The tumor derived from CD44-high and low cells were collected at endpoint and FACS was performed to measure CD44 status between CD44hi and CD44low tumors. **H.** CD44 expression in CD44-high derived tumor (**left**) and CD44-low derived tumor (**right**) by IF staining.

### Iron is key regulator in CD44hi subpopulation

To investigate the tumorigenic potential of the CD44hi subpopulation, we performed RNA-seq on FACS-sorted CD44hi and CD44low cells from LAPC9-GFP tumors. Transcriptomic profiling revealed distinct expression programs between the two subpopulations, confirming that CD44hi cells harbor unique molecular features (Supplementary Fig. 2A). Pathway enrichment analysis demonstrated significant upregulation of MYC targets, cholesterol homeostasis, and fatty acid metabolism in CD44hi cells (Supplementary Fig. 2B). CD44hi cells exhibited an SCL-like signature, whereas CD44low cells were enriched for AR-related features (Figure 3A; Supplementary Fig. 2C), consistent with prior report designating CD44 as a marker of the CRPC-SCL subtype [4]. Critically, iron metabolism pathways were also enriched, and CD44hi cells displayed elevated iron-related features (Figure 3B). Previous studies have shown that CD44 mediates iron uptake via HA [7]. In line with this, HA treatment increased intracellular iron levels in the CD44hi model (Supplementary Fig. 2D-E). Within LAPC9 tumors, the CD44hi subpopulation exhibited significantly higher intracellular iron levels compared with CD44low cells (Figure 3C). Consistently, western blot analysis revealed upregulation of iron -associated proteins, including Nuclear factor erythroid 2-related factor 2 (NRF2) and Ferritin Light Chain (FTL), in CD44hi cells (Figure 3D). To validate these findings *in vitro*, we utilized PC3 cells, a CRPC-SCL subtype characterized by high CD44 expression [4]. In order to mimic CD44hi and CD44low status in cell line, CD44 expression was silenced in PC3 cells to generate PC3-shCD44 cells, enabling comparison between CD44 high and CD44 deficient states. Similarly, as the 22Rv1 cell line exhibits heterogeneous CD44 expression (Supplementary Fig. 2F), FACS was used to isolate CD44hi and CD44low cells, establishing 22RV1-CD44hi and 22RV1-CD44low cell lines (Supplementary Fig. 2G-H). Measurement of intracellular iron levels revealed that CD44hi cells maintained elevated iron content relative to their CD44-knockdown counterparts (Figure 3E-F), corroborating the findings observed in the LAPC9 model. Together, these findings establish that CD44hi cells exhibit elevated intracellular iron compared with CD44low cells. We next asked whether iron is functionally required for sustaining CD44 expression. Treatment of 22Rv1-CD44hi cells with the iron chelator deferoxamine (DFO) reduced CD44 expression (Figure 3G). Given that CD44low cells re-expression CD44 *in vivo* during tumor growth (Figure 2G), we next investigated whether iron availability regulates this process. In order to mimic this CD44 conversion *in vitro*, the 22RV1-CD44low cells were cultured in 3D condition, the 22Rv1-CD44low cells exhibited a progressive increase in CD44 expression, indicative of CD44low to CD44hi conversion (Figure. 3H). Notably, treatment with the DFO effectively blocked this transition (Figure. 3H). Together, these findings demonstrate that iron is essential for maintaining CD44 expression and promoting CD44 conversion. Given that iron activates JMJD domain-containing demethylases, we examined histone demethylases expression in CD44 subpopulations in RNA-seq data. The H3K9me2-associated demethylases, like KDM3A and KDM3B, were upregulated in CD44hi cells (Figure 3I). As H3K9me2 is a repressive chromatin mark, we compared its levels across subpopulations and found H3K9me2 enrichment in LAPC9 CD44low cells (Figure 3J, Supplementary Fig. 3A). In cell line models, H3K9me2 levels were enriched in CD44low or shCD44 cells (Figure 3K-L). Collectively, these results indicate that CD44hi cells maintain elevated intracellular iron levels, whereas H3K9me2 modification is preferentially enriched in CD44low cells.

**Fig 3.**
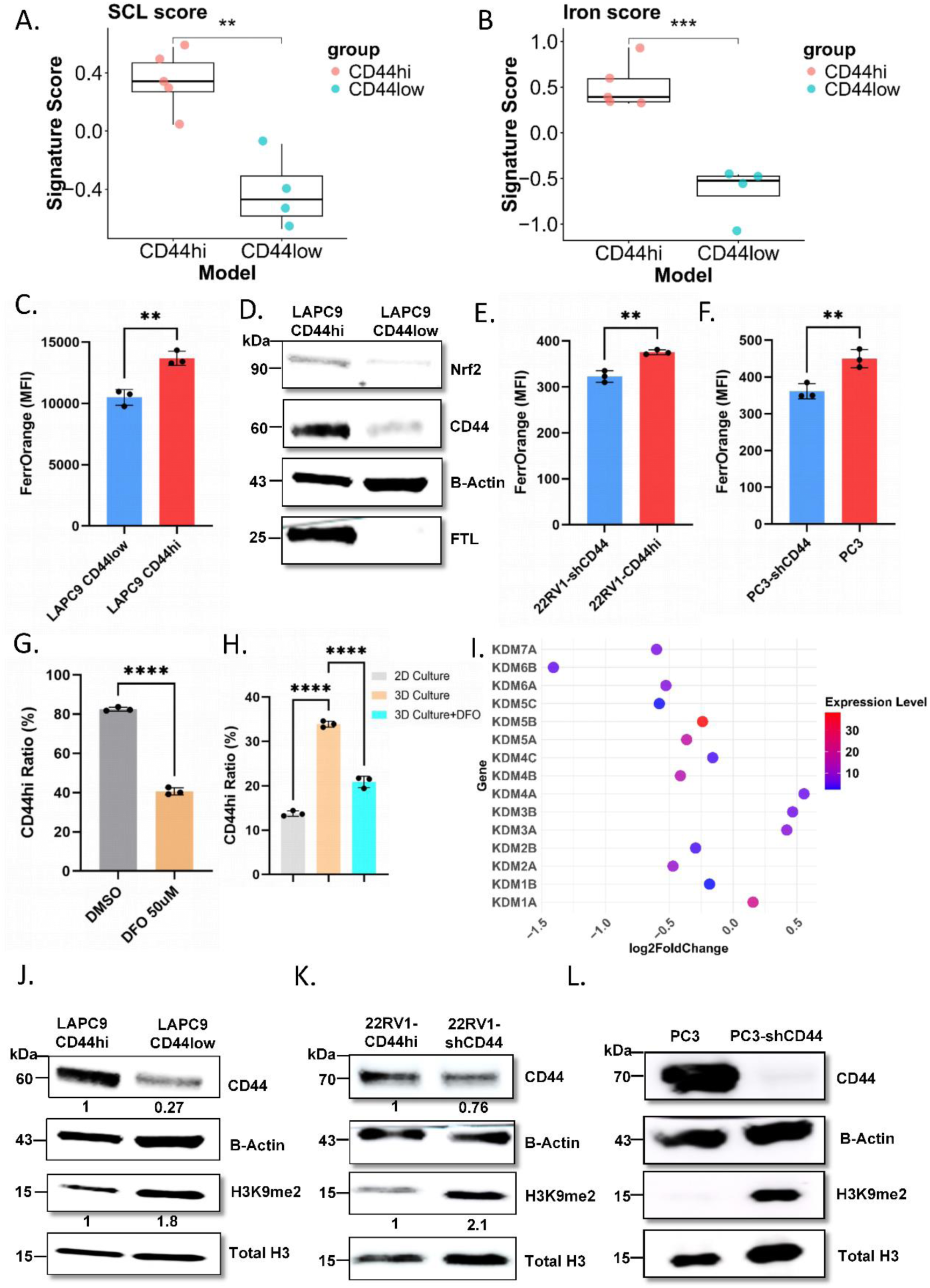
RNA-seq analysis on LAPC9 CD44hi and CD44low subpopulations reveals Iron as key regulator. **A-B.** identified significant pathways in LAPC9 CD44hi subpopulation. Signature analysis was identified stem cell like (SCL) score (**A**) and iron score (**B**) between CD44hi and CD44low cells. **C.** FACS performed on LAPC9-GFP tumor cells to sort CD44hi from CD44low cells and FerrOrange to reveal intracellular iron levels (n=3). **D.** Protein levels of CD44, NRF2 and FTL in the sorted subpopulations. The b-Actin was used as loading control. **E.** Intracellular iron levels on stably expressing 22RV1-CD44hi and 22RV1-shCD44 KD cells stained with FerrOrange (n=3). **F.** Intracellular iron levels on stably expressing PC3 and PC3-shCD44 KD cells stained with FerrOrange (n=3). **G.** FACS staining with CD44-APC were performed on 22RV1 cells treated with or without DFO (50uM) for 72h (n=3). Ratio of CD44hi cells were compared between two groups. **H.** 22RV1-CD44low cells cultured in 2D, 3D organoid medium condition and with treat DFO 30uM for 3 days. FACS staining with CD44-APC. Ratio of CD44hi cells were compared among different groups (n=3). **I.** The expression level of demethylases between LAPC9 CD44hi and CD44low subpopulation based on RNA-seq data. **J-L.** Western Blot for CD44 and H3K9me2 level between sorted CD44hi and CD44low cells from LAPC9-GFP tumor (**J**), between 22RV1-CD44hi and 22RV1-shCD44 cells (**K**). and between PC3 and PC3-shCD44 cells (**L**).

### Iron regulate CD44 expression through H3K9me2 demethylase KDM3A

To directly test whether H3K9me2 regulates CD44 transcription, we performed ChIP followed by qPCR (ChIP-qPCR) to assess H3K9me2 mark at the CD44 locus. H3K9me2 was specifically enriched at the CD44 promoter region (Figure 4A), consistent with its role as a transcriptional repressive modification [15]. Comparison of 22Rv1-CD44hi and CD44low cells revealed higher H3K9me2 occupancy at the CD44 locus in CD44low cells (Figure 4B), which corresponded to reduced CD44 transcript levels relative to CD44hi cells (Figure 4C). We next investigated whether iron regulates CD44 expression through H3K9me2 modification. Treatment of 22Rv1-CD44hi cells with the DFO markedly reduced CD44 expression at both the mRNA and protein levels, while concomitantly increasing H3K9me2 modification (Figure 4D-E). Given that iron can activate Jumonji-domain histone demethylases [9], we next treated 22Rv1-CD44hi cells with the pan-H3K9 demethylase inhibitor IOX1 [16]. IOX1 treatment decreased CD44 expression at both transcript and protein levels (Supplementary Fig. 3B-C) and prevented the transition of CD44low cells into the CD44hi state (Supplementary Fig. 3D), supporting a role for H3K9 demethylases in regulating CD44 expression. Based on RNA-seq analysis, KDM3A, an H3K9me2-specific demethylase [17], was found upregulated in CD44hi cells, highlighting its putative role in modulating CD44 expression (Figure 3I). Pharmacological inhibition of KDM3A using the selective inhibitor CBA-1 [18] reduced CD44 expression and increased H3K9me2 levels (Figure 4F-G). Furthermore, during the CD44low to CD44hi conversion process, H3K9me2 levels declined, whereas treatment with CBA-1 restored H3K9me2 and blocked CD44 induction (Figure 4H-I). Together, these findings support a model in which iron sustains CD44 expression by activating KDM3A-mediated removal of repressive H3K9me2 marks at the CD44 locus.

**Figure. 4.**
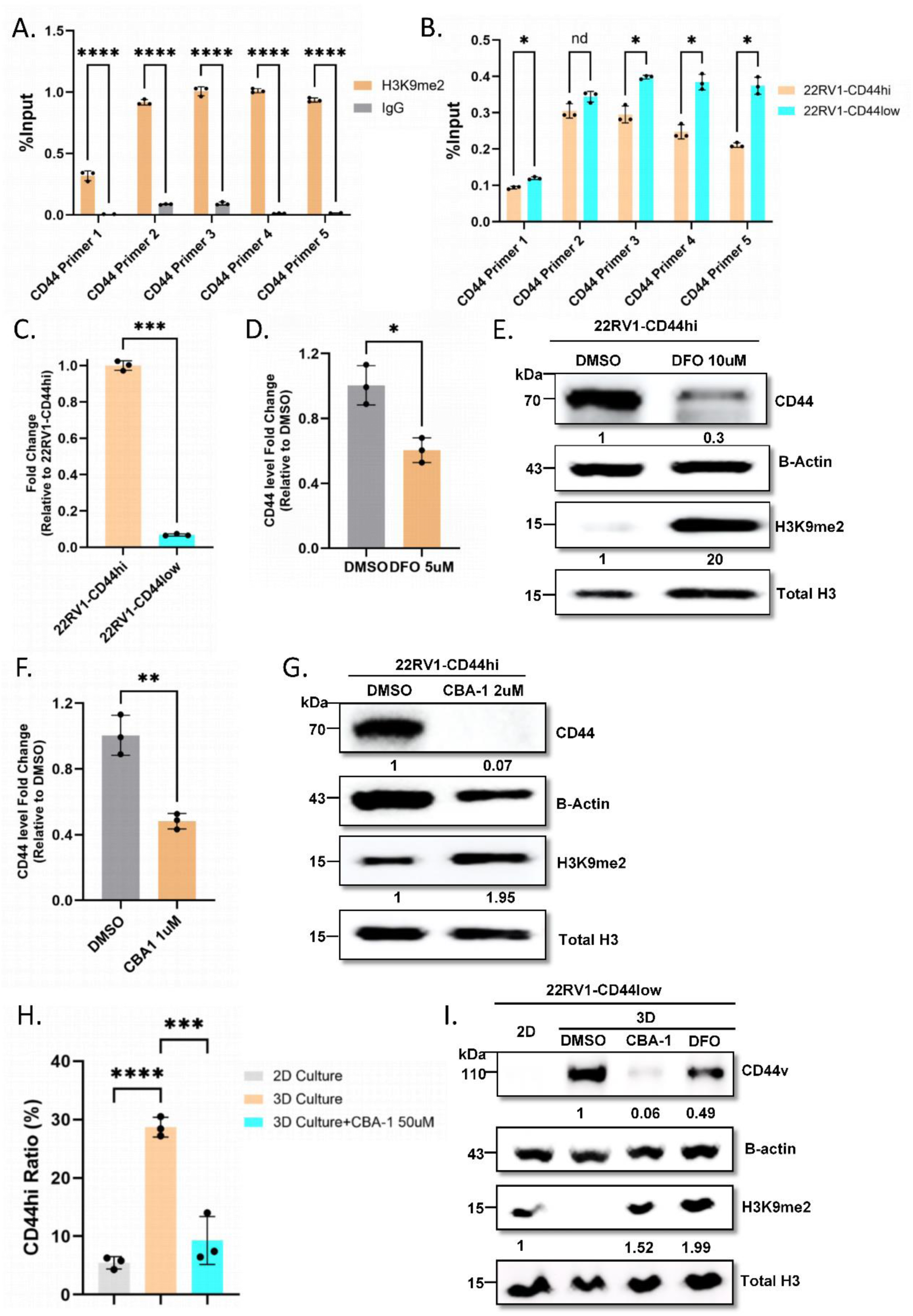
Iron levels directly mediate CD44 status conversion. **A.** Comparing CD44 between H3K9me2 and IgG chromatin immunoprecipitation in 22Rv1-CD44hi cells by Chip-qPCR (n=3). The CD44 primers were designed based on the upstream region of transcriptional factor bind sit (from 0-400bp). **B.** Chip-qPCR compare CD44 between 22RV1-CD44hi cells and 22RV1-CD44low cells (n=3). **C.** qPCR comparing CD44 between 22RV1-CD44hi cells and treated with 5uM DFO for 72h (n=3). **D.** qPCR compare CD44 between 22RV1-CD44hi and treated with 10uM DFO for 72h (n=3). **E.** Western Blot for CD44 and H3K9me2 level between 22RV1-CD44hi and treated with 10uM DFO for 72h. **F.** qPCR compare CD44 between 22RV1-CD44hi cells and treated with 1uM CBA-1 for 72h. **G.** Western Blot for CD44 and H3K9me2 level between 22RV1-CD44hi and treated with 2uM CBA-1 for 72h. **H.** 22RV1-CD44low cells cultured in 2D, 3D organoid medium condition and with treat CBA-1 50uM for 3 days (n=3). **I.** Western Blot for 22RV1-CD44low cells cultured in 2D, 3D organoid medium condition and with treat CBA-150uM and DFO 30uM 3 days.

### Iron regulates features of CD44hi cells by modulating H3K9me2

We next asked whether iron also impact the functional properties of CD44hi cells. CD44hi cells demonstrated significantly greater clonogenic capacity compared to shCD44 cells (Figure 5A-D), consistent with our previous findings in LAPC9 models (Figure 2B, E). To assess the role of iron in maintaining these phenotypic features, CD44hi cells were treated with the iron chelator DFO. Iron depletion markedly reduced the colony-forming ability and proliferation of CD44hi cells (Figure 5E-G, Supplementary Fig. 3E-G). As iron depletion increases H3K9me2 mark, thereby repressing gene expression (Figure 4E), we next examined whether H3K9me2 regulation contributes to the tumorigenic properties of CD44hi cells. Pharmacological inhibition of KDM3A similarly suppressed colony formation and proliferation of CD44hi cells (Figure 5H-J, Supplementary Fig. 3H-J). Together, these data demonstrate that iron sustains the tumorigenic properties of CD44hi cells through KDM3A-mediated regulation of H3K9me2, highlighting iron metabolism as a critical determinant of the CD44hi state.

**Figure 5.**
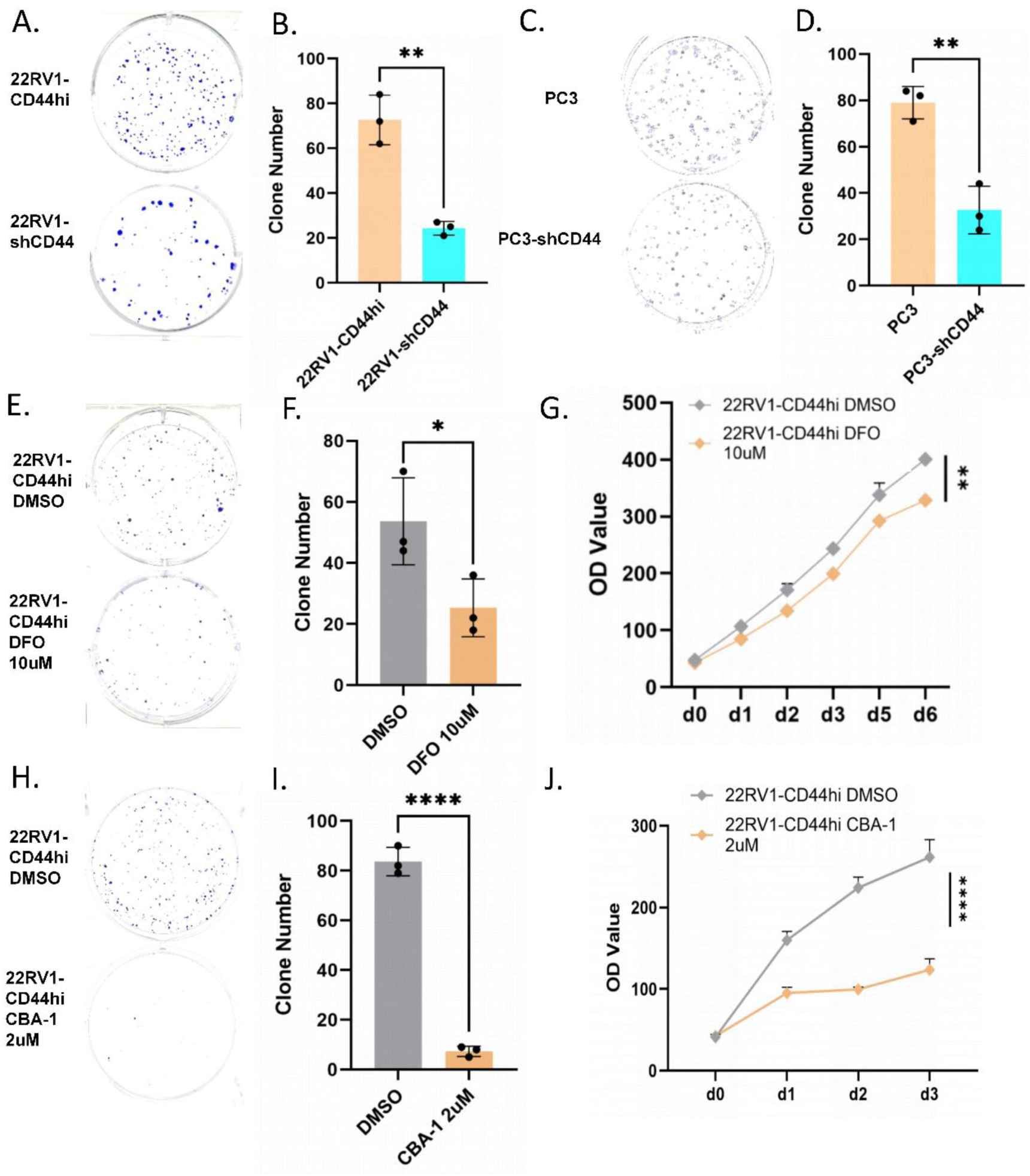
Iron regulate features of CD44hi cells by modulating H3K9me2. **A.** Clonogenic assay of 22Rv1-CD44hi cells and 22Rv1-shCD44 cells (1000 cells/well, n=3). **B.** Measured and quantified the clone number by ImageJ. **C.** Clonogenic assay of PC3 cells and PC3-shCD44 cells (500 cells/well, n=3). **D.** Measured and quantified the clone number by ImageJ. **E.** Clonogenic assay of 22RV1-CD44hi cells treated with DMSO or DFO 10uM for 72 h before seeding (1000 cells/well, n=3). **F.** Measured and quantified the clone number by ImageJ. **G.** Measurement growth curve of 22RV1-CD44hi treated with DMSO or DFO 10uM, viability measured by PrestoBlue (n=4). **H.** Clonogenic assay of 22RV1-CD44hi cells treated with DMSO or CBSA-1 2uM for 72 h before seeding (1000 cells/well, n=3). **I.** Measured and quantified the clone number by ImageJ. **J.** Measurement growth curve of 22RV1-CD44hi treated with DMSO or CBA-1 2uM, viability measured by PrestoBlue (n=5).

### NRF2 is highly active in CD44hi cells, and its inhibition increases their sensitivity to cell death

Since CD44hi cells represent an aggressive subpopulation within CRPC-SCL tumors and maintain elevated intracellular iron, we next sought strategies to specifically target this subpopulation. Analysis of our RNA-seq data and transcriptional regulatory networks identified NRF2, a master regulator of antioxidant and ferroptosis pathways, as highly active in CD44hi cells (Figure 6A). NRF2 activation was also observed in 22Rv1 and PC3 cell lines (Figure 6B-C; Supplementary Fig.4B). Given that iron promotes reactive oxygen species (ROS) and NRF2 is a central antioxidant regulator [19], we tested whether iron directly activates NRF2 in CD44hi cells. Supplementation of iron in 22RV1-shCD44 cells increased NRF2 activity, which was reversed by DFO treatment (Figure 6D-E). Consistently, iron depletion with DFO reduced NRF2 activity in 22Rv1-CD44hi cells (Supplementary Fig. 4C-D). The same result also observed in PC3 cell line (Supplementary Fig. 4E). These results indicate that CD44hi cells possess elevated intercellular iron and enhanced NRF2 activity. To functionally assess the therapeutic vulnerability of this pathway, we treated cells with the NRF2 inhibitor Brusatol [20]. CD44hi cells were more sensitive to Brusatol than CD44low cells (Figure 6F-G). In a competitive assay, mixed cultures of CD44hi and shCD44-GFP cells revealed preferential elimination of CD44hi cells upon Brusatol treatment at equivalent concentrations (Figure 6H-I, Supplementary Fig.4G-I). *Ex vivo* treatment of LAPC9 tumor tissues further confirmed that Brusatol selectively reduced the CD44hi subpopulation (Figure 6J-K). Together, these data demonstrate that NRF2 is activated by iron in CD44hi cells and that pharmacological inhibition of NRF2 preferentially targets this aggressive CRPC-SCL subpopulation.

**Fig 6.**
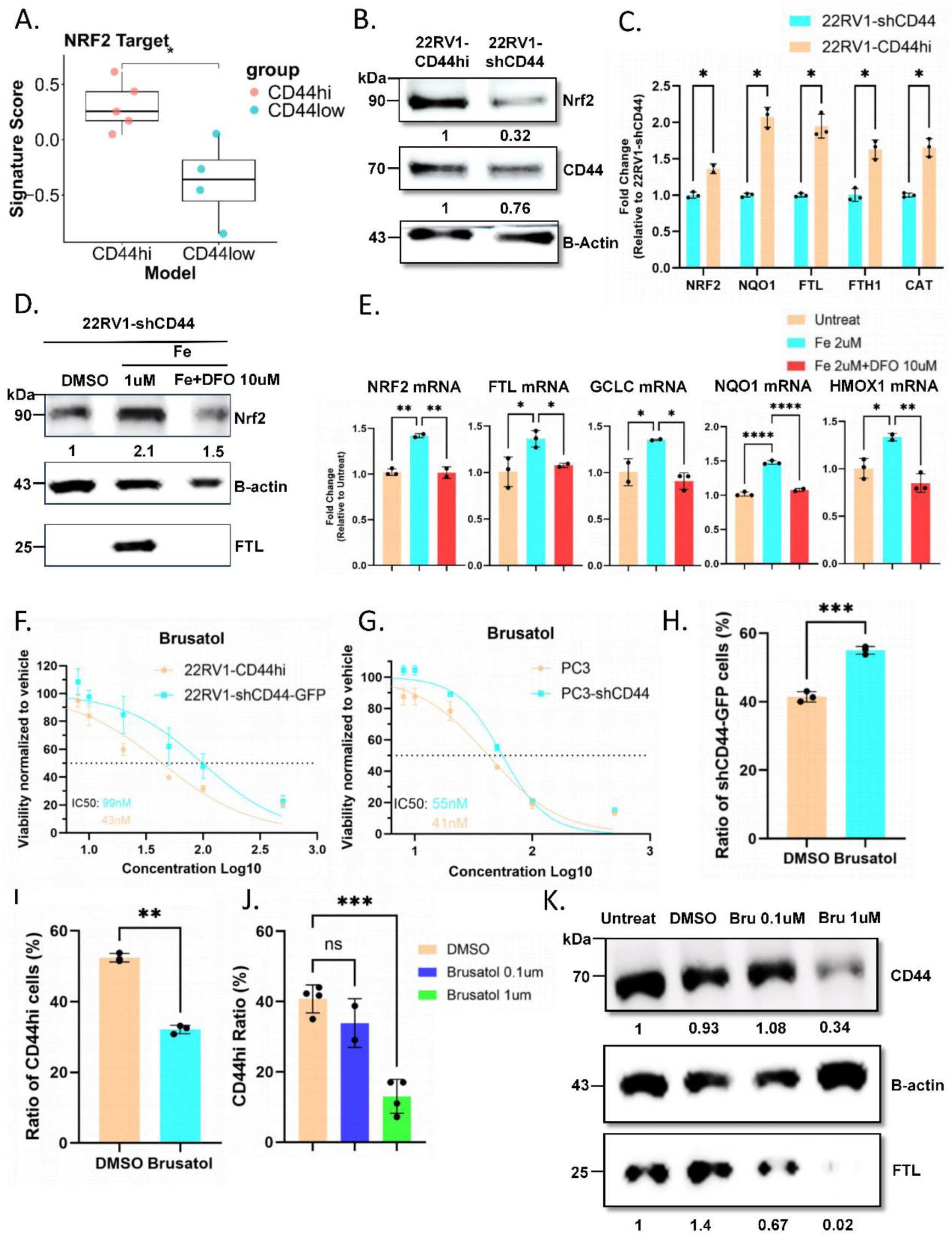
Iron levels regulate NRF2 activity in CD44hi cells and NRF2 inhibition increases the sensitivity of CD44hi cells to cell death. **A.** Signature analysis was identified NRF2 target genes level between CD44hi and CD44low cells. **B.** Western-blot confirming CD44 and NRF2 level in 22RV1-CD44hi and 22RV1-shCD44 cells. The b-Actin was used as loading control. Signal of protein were quantified by Image J. **C.** qPCR for mRNA level of NRF2 downstream genes between 22RV1-CD44hi and 22RV1-shCD44 cells (n=3). HRPT and β-actin were used as housekeeping genes. unpaired t-test for CD44hi vs shCD44, p< 0.05 (*), p< 0.005 (**), p< 0.001 (***), p< 0.0001 (****). **D.** Western Blotting analysis of NRF2 and FTL levels. Protein detection signals were quantified by ImageJ. 22RV1-shCD44 cells treated with iron (1uM) and iron chelator DFO (5uM) for 72h. **E.** qPCR for mRNA level of NRF2 downstream genes. 22RV1-shCD44 cells treated with Iron (2uM)+DFO (10uM) for 72h (n=3). One-way Anova for E, p< 0.05 (*), p< 0.005 (**), p< 0.001 (***), p< 0.0001 (****). **F.** Measurement of 22RV1-CD44hi and 22RV1-shCD44 cells viability by PrestoBlue, Brusatol treated cells for 72h (n=6). **E.** Measurement of PC3and PC3-shCD44 cells viability by PrestoBlue, Brusatol treated cells for 72h (n=6). **H-I.** Mix 22RV1-CD44hi cells and 22RV1-shCD44-GFP cells as 1:1 ratio, treated with 0.05uM Brusatol for 72h. Preformed FACS check ratio of CD44hi and CD44low cells with DMSO group. The GFP+ ratio (**H**) and CD44hi ratio (**I**) were compared by chi-square test (n=3). **J, K.** LAPC9 fresh tissue was cut by 2mm slice and culture as ex vivo. Treated Brusatol with (01uM and 1uM) for 4 days, collect tissue for FACS (**J**) by staining CD44-APC and for WB (**K**).

### Brusatol induces Ferroptosis in CD44hi cells through NRF2 inhibition

NRF2 serves as a major antioxidant regulator and has been shown to influence iron metabolism by suppressing ferritinophagy, the autophagic degradation of ferritin that releases Fe^2+^, thereby preventing excessive accumulation of the labile iron pool (LIP) [21]. Based on this, we hypothesized that NRF2 inhibition would further increase LIP in CD44hi cells. Indeed, treatment of CD44hi cells with the NRF2 inhibitor Brusatol reduced NRF2 activity (Figure 7A-B, Supplementary Fig.5A). Moreover, treatment with Brusatol led to a greater increase in LIP levels in CD44hi cells compared to shCD44 cells (Figure 7C-D, Supplementary Fig.5B-C). Because LIP promotes lipid peroxidation and ferroptosis [22], we next examined whether NRF2 inhibition sensitizes CD44hi cells to ferroptotic stress. At baseline, CD44hi cells exhibited higher lipid peroxidation than shCD44 controls. Brusatol treatment markedly enhanced lipid peroxidation in CD44hi cells compared to shCD44 cells, which was reversed by either an iron chelator (DFO) or a lipid peroxidation inhibitor (liproxstatin-1) (Figure 7E-F, Supplementary Fig. 5D-E). Functionally, Brusatol reduced the viability of CD44hi cells more strongly than shCD44 cells, and this effect was rescued by ferroptosis inhibitors (Figure 7G, Supplementary Fig. 5F), confirming ferroptosis as the primary mode of cell death. To further validate this, we tested whether other cell death pathways contributed to Brusatol-induced cytotoxicity. CD44hi cells were treated with Brusatol in combination with inhibitors of apoptosis (ZVAD-FMK), autophagy (chloroquine, CQ), or ferroptosis (liproxstatin-1). Only DFO rescued Brusatol-induced cell death, whereas apoptosis and autophagy inhibition had minimal effects (Figure 7H). Together, these results demonstrate that CD44hi cells are uniquely vulnerable to NRF2 inhibition due to their elevated iron content. Blocking NRF2 increases LIP and drives ferroptosis selectively in CD44hi cells, establishing ferroptosis induction as a therapeutic strategy for targeting this aggressive CRPC-SCL subpopulation.

**Figure. 7.**
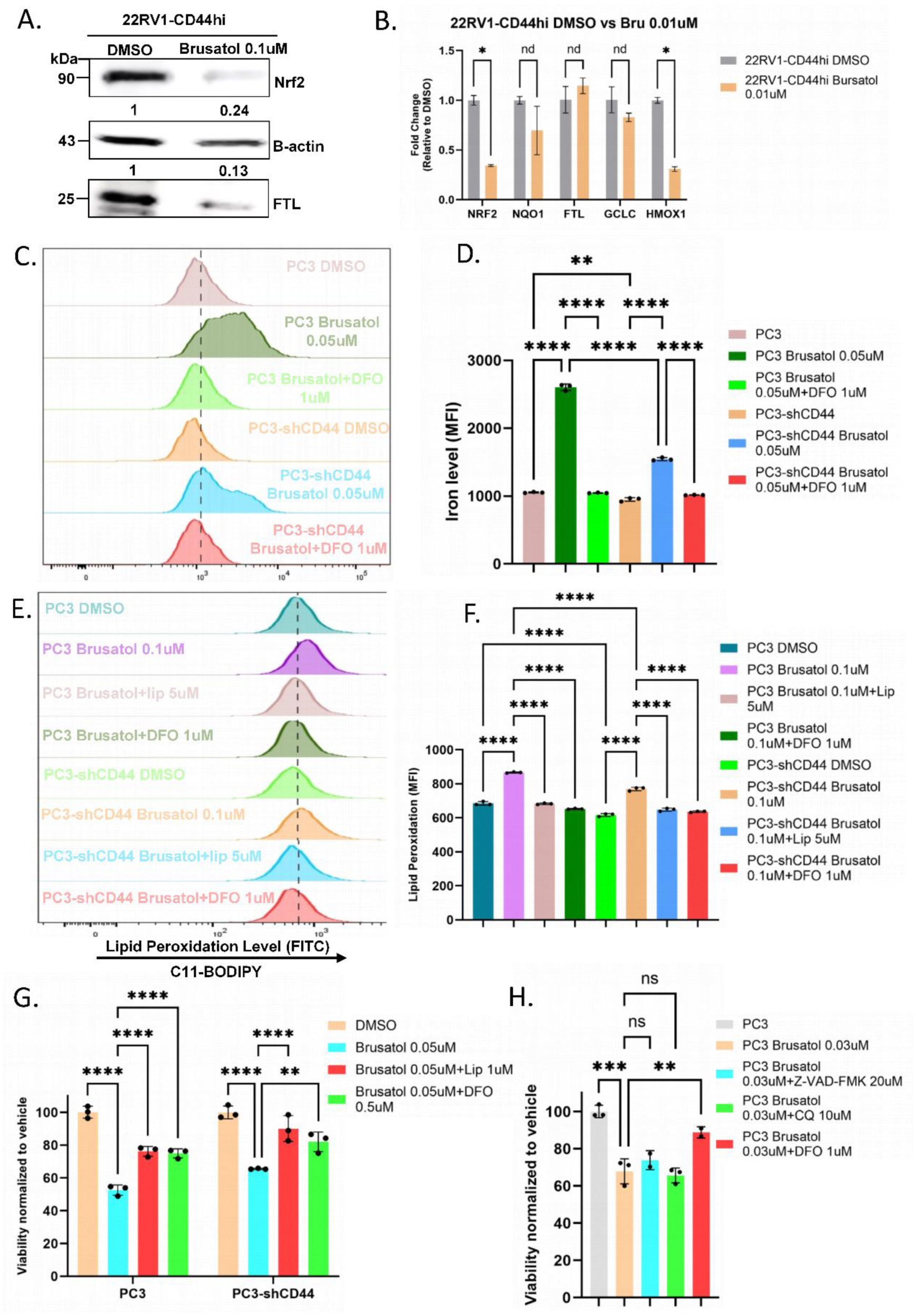
Brusatol induces Ferroptosis in CD44hi cells through NRF2 inhibition. **A.** Western Blotting analysis for NRF2 and FTL protein levels. 22RV1-CD44hi cells treated with Brusatol (0.1uM) for 72h. Protein detection signals were quantified by ImageJ. B. qPCR for mRNA level of NRF2 downstream genes (n=3). **C, D.** Labile iron pool level after Brusatol treatment. FACS (staining with FerrOrange) on PC3 cells and PC3-shCD44 cells treated with Brusatol (0.05uM) with or without DFO (1uM), for 72h (**C**), iron levels were compared among six groups (n=3) (**D**). **E, F.** Lipid peroxidation levels after Brusatol treatment. FACS (staining with C11-BODIPY) on PC3 and PC3-shCD44 cells treated with Brusatol (0.1uM) with or without Lipostatin-1 (5uM) and DFO (1uM) for 24h (**E**), lipid peroxidation levels were compared among different groups (n=3) (**F**). **G.** Measurement of PC3 and PC3-shCD44 cells viability by PrestoBlue, Brusatol (0.05uM) treated cells with or without Lipostatin-1 (1uM) and DFO (0.5uM) for 72h (n=3). **H.** Measurement of PC3 cells viability by PrestoBlue, Brusatol (0.03uM) treated cells with or without apoptosis inhibitor Z-VAD-FMK (20uM), Autophagy inhibitor CQ (10uM) and ferroptosis inhibitor DFO (1uM) for 72h (n=3).

### Brusatol inhibit CRPC-SCL tumor growth *in vivo*

To evaluate the impact of NRF2 inhibition on tumor growth *in vivo*, we established subcutaneous LAPC9-copGFP-CBR xenografts in CB17 SCID mice. At one-week post-implantation, mice were randomized by bioluminescence intensity and body weight and subsequently treated with DMSO or Brusatol (2 mg/kg every other day) for two weeks (Figure 8A). Bioluminescence imaging at week 3 revealed significantly reduced tumor signal in the Brusatol treated group compared to vehicle-treated controls (Figure 8B). Consistently, endpoint tumor weights were lower in the Brusatol treated group (Figure 8C-D), indicating that NRF2 inhibition suppresses CRPC-SCL tumor growth *in vivo*. We next assessed whether Brusatol selectively targets CD44hi cells within tumors. Flow cytometry analysis demonstrated a marked reduction in CD44hi cells in Brusatol-treated tumors compared with controls (Figure 8E). NRF2 and ferritin levels were assessed in the DMSO and Brusatol treated groups to verify the inhibitory effect of Brusatol on NRF2 signaling. A reduction in CD44 expression in the Brusatol treated group was further confirmed by FACS analysis. (Figure 8F). Immunohistochemical staining demonstrated that the expression levels of CD44 and NRF2 were reduced following Brusatol treatment. Moreover, the lipid peroxidation marker 4-hydroxynonenal (4-HNE) was increased in treatment group indicating the tumor underwent with ferroptosis after Brusatol treatment (Figure 8G). This finding is consistent with our *ex vivo* observations (Figure 6J-K), and supports the conclusion that NRF2 inhibition specifically diminishes the CD44hi subpopulation. Together, these results demonstrate that pharmacological inhibition of NRF2 impairs tumor growth and reduces CD44hi cells in CRPC-SCL tumors, highlighting NRF2 as a therapeutic vulnerability in this aggressive prostate cancer subtype.

**Figure 8.**
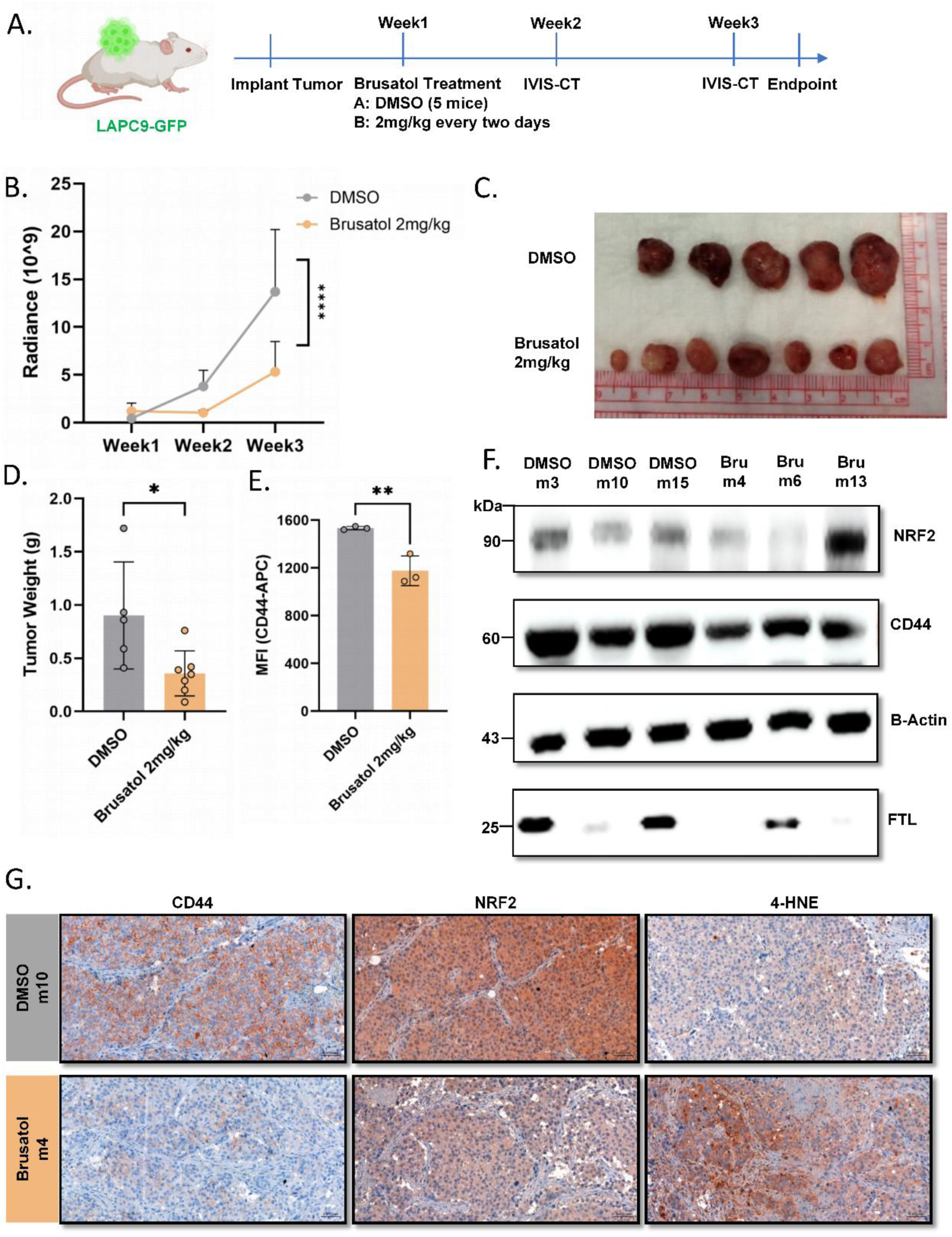
Brusatol inhibit CRPC-SCL tumor growth in vivo. **A**. Scheme for Brusatol treatment in vivo, brusatol was IP inject 0.25mg/kg every day or 2mg/kg every two days (5 mice/group). **B.** Luciferase was measured by IVIS-CT in DMSO and Brusatol 2mg/kg groups weekly. The quantified radiance was used to measure tumor dynamics in each group. **C.** The tumors were collected at endpoint, tumor sizes were measured by scale (Hight*Length*Width*0.8), tumor weight were measured by scale (g). unpaired t-test for each group (5 mice/group). **D, E.** FACS was performed to measure CD44 status between DMSO and Brusatol 2mg/kg tumors (**D**). Median Fluorescence Intensity (MFI) were compared between two groups (n=3) by unpaired t-test (**E**). **F.** Western-blot were performed on DMSO and Brusatol 2mg/kg tumors to measure NRF2 and FTL level. **G.** Immunohistochemical staining were performed on DMSO and Brusatol 2mg/kg tumors to measure NRF2, CD44 and 4-HNE level.

### Enrichment of CD44 and iron-related genes associated with unfavorable clinical outcome in PCa

To evaluate NRF2 as a therapeutic vulnerability in CRPC-SCL tumors, we treated multiple PCa organoid models with Brusatol, including LAPC9 (SCL subtype), PM154 (NE subtype), PCa16 (Wnt-driven subtype), and BM18 (androgen-dependent subtype). LAPC9 organoids exhibited greater sensitivity to Brusatol compared to the other PCa subtypes (Figure 9A). These findings indicate that Brusatol preferentially targets CD44hi cells with SCL features and suggest that NRF2 inhibition may represent a potential therapeutic strategy for CRPC-SCL tumors.

**Figure. 9.**
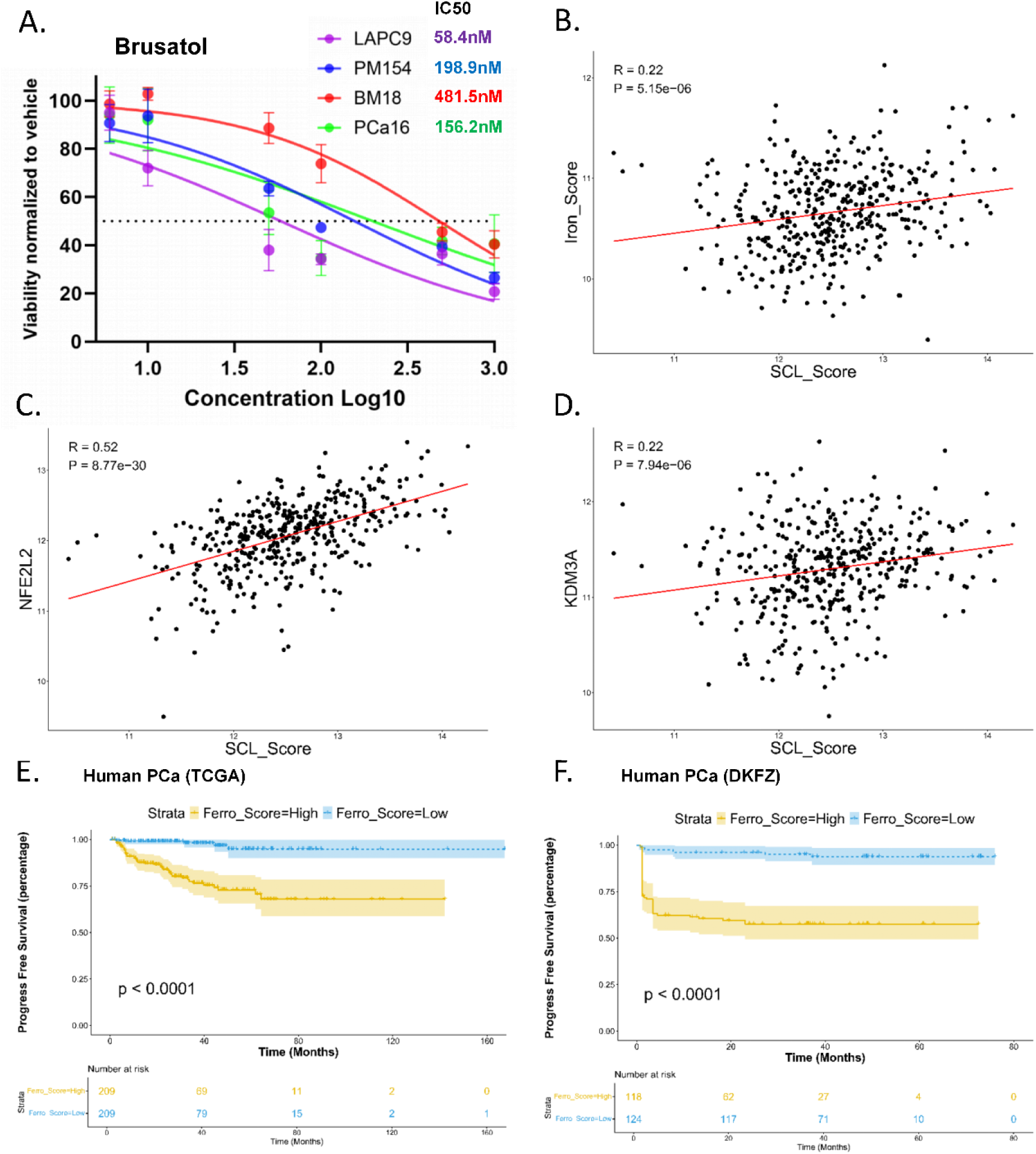
CD44 and iron abundance in human PCa predict unfavorable clinical outcomes. **A.** Brusatol treatment on different type PCa organoids (AR, Wnt, NE and SCL-like) for 48h, SCL tumor LAPC9 was sensitive to Brusatol. **B-D.** Pearson correlation analyses reveal positive associations of stem cell-like (SCL) Score with iron score (**B**), NFE2L2 (NRF2) (**C**) and KDM3A level (**D**). This dataset of TCGA human PCa contains 418 patients (n = 418 patients). **E-F.** Using TCGA bulk RNA-seq (TCGA, n = 418 patients, **E**) and a DKFZdataset (n = 242 patients **F**), the FerroScore high PCa patients exhibit a significantly higher biochemical recurrence (BCR) rate as compared to FerroScore-low subjects. Kaplan–Meier (K-M) analyses of BCR based on FerroScore.

**Figure. 10.**
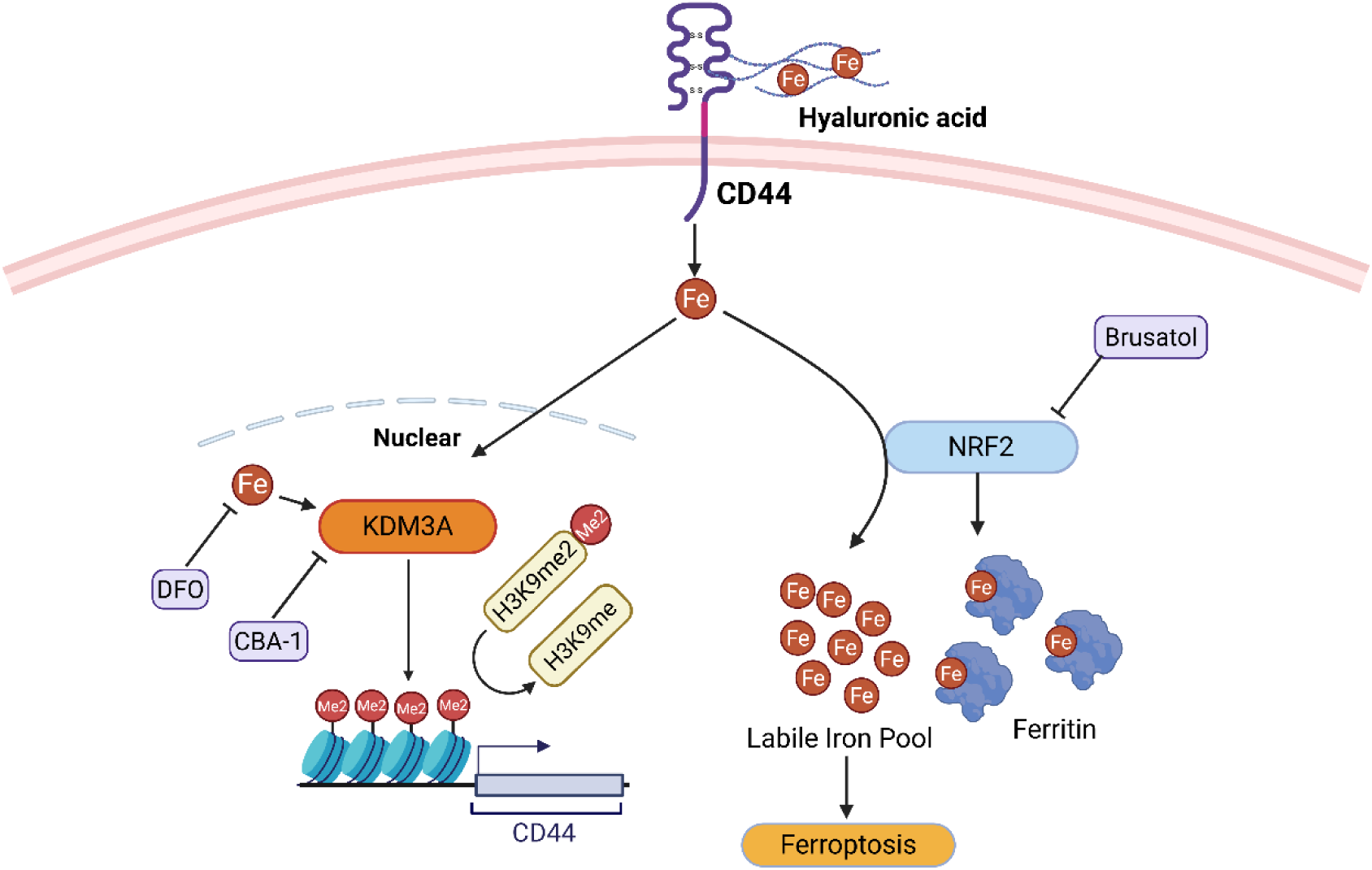
Iron as a key regulator of CD44 expression and SCL characteristics through H3K9me2 modification. CD44hi cells exhibit enhanced iron uptake, which facilitates nuclear activation of the H3K9me2 demethylase KDM3A. Activated KDM3A promotes CD44 expression and sustains stem cell–like properties. Elevated intracellular iron levels upregulate NRF2 and ferritin, contributing to iron homeostasis by stabilizing free iron. Pharmacological inhibition of NRF2 with Brusatol reduces ferritin expression, thereby releasing stored iron and increasing the labile iron pool, ultimately sensitizing cells to ferroptosis.

To extend these findings to patients, we assessed the relationship between SCL biology and iron-linked regulation in TCGA bulk RNA-seq data. The SCL score positively correlated with the iron score, KDM3A expression, and NRF2 (NFE2L2) levels (Figure 9B-D), aligning with our mechanistic data. We then defined a CD44hi-informed gene set (*CD44*, *FTL*, *FTH1*, *SLC11A2*, *KDM3A*, and *NFE2L2*; “FerroScore”) and found that high FerroScore was associated with a higher rate of biochemical recurrence (BCR) and significantly shorter progression-free survival in TCGA and DKFZ cohorts (Fig. 9E, F). Collectively, these data link enrichment of CD44 and iron-regulated programs to adverse clinical outcomes and nominate the CD44-iron-NRF2 axis as a clinically relevant target in PCa.

## Discussion

In this study, we focused on the CD44hi subpopulation within the CRPC-SCL subtype. We found that CD44hi cells display SCL-associated transcriptional features and exhibit greater tumorigenic potential *in vitro* and *in vivo* compared with CD44low cells. Mechanistically, CD44hi cells maintain elevated intracellular iron, which shapes their identity by regulating CD44 expression through iron-dependent epigenetic mechanisms. Leveraging this vulnerability, we demonstrated that inhibition of NRF2, a key antioxidant regulator, further increases labile iron and triggers ferroptosis, thereby selectively targeting CD44hi cells.

Recent chromatin profiling studies have classified CRPC into distinct molecular subtypes, including androgen receptor (AR)-driven, Wnt-driven, neuroendocrine (NE), and stem cell-like (SCL) subtypes [4]. Among these, the CRPC-SCL subtype represents a newly defined category characterized by enrichment of a mammary stem cell gene signature and elevated expression of cancer stem cell markers, such as CD44 and TROP2A. In this study, we report for the first time that CD44hi tumor cells constitute a distinct subpopulation within CRPC and exhibit molecular and functional features consistent with the SCL subtype. Our *in vivo* data demonstrate that CD44hi cells represent a more aggressive subpopulation within CRPC-SCL tumors, highlighting a critical role for CD44 in driving SCL tumor characteristics. Moreover, consistent with previous findings [23], CD44low cells tend to convert to a CD44hi state during tumor growth, suggesting that CRPC-SCL tumors preferentially maintain CD44 expression to preserve their SCL characteristics. CD44 has previously been identified as a cancer stem cell marker in primary prostate cancer [6]. According to the clonal progression model, CRPC arises from specific subclones that survive androgen deprivation and subsequently expand following castration. Our findings support the notion that CD44hi cells represent such a subpopulation, capable of persisting under androgen-deprived conditions and driving tumor outgrowth within the CRPC-SCL subtype. These observations raise the possibility that CRPC-SCL tumors may originate from CD44hi progenitor cells. Future lineage-tracing studies will be necessary to elucidate the evolutionary dynamics and fate of CD44hi cells during CRPC progression.

Iron plays a central role in epigenetic regulation, and recent studies have shown that CD44 sustains high intracellular iron, cooperating to regulate epithelial–mesenchymal transition in CD44+ cancer cells and inflammatory responses in macrophages [24].

Consistently, we confirmed that CD44hi cells in CRPC-SCL tumors maintain elevated iron levels, which in turn shape their phenotypic and transcriptional features. Mechanistically, iron modulates H3K9 methylation in CD44hi cells, as iron chelation led to increased H3K9me2 levels. Previous studies [25,26] have implicated key transcriptional regulators in CD44hi cells, including MYC, SREBP1, and CEBPA, in H3K9me2-mediated chromatin regulation, suggesting a potential link between iron availability and the maintenance of CD44hi cell identity to support tumor growth. Given that iron positively regulates CD44 expression, the interplay between CD44 and H3K9me2 may serve as a useful marker for defining iron-loaded subpopulations or molecular subtypes in prostate cancer. However, iron functions as a broad regulator of epigenetic processes. Although our data demonstrate that modulation of H3K9me2 suppresses clonogenic capacity and cellular proliferation, chromatin profiling comparing CD44hi and CD44low cells are required to comprehensively delineate the epigenetic landscape of CD44hi cells and to better define chromatin accessibility changes associated with this subpopulation.

To exploit the vulnerability conferred by elevated iron levels in CD44hi cells, we examined the role of NRF2, a key regulator of oxidative stress responses. We observed that NRF2 expression was markedly upregulated in CD44hi cells, consistent with previous findings in breast cancer, where P62-mediated regulation enhanced NRF2 activity [27]. In our study, we further establish a link between intracellular iron levels and NRF2 activation. In agreement with a previous report [28], we found that increased iron promotes NRF2 expression and activates its downstream target genes. In this study, we demonstrate that NRF2 regulates iron metabolism, and that inhibition of NRF2 leads to an increase in the LIP. Similar findings have been reported in NRF2-deficient cells, where loss of NRF2 led to increased NCOA4 expression, enhanced recruitment of apoferritin into autophagosomes, and disrupted ferritinophagy, collectively resulting in elevated LIP and heightened ferroptosis sensitivity [21]. Moreover, pharmacological inhibition of NRF2 using Brusatol has been shown to induce ferroptosis by depleting glutathione (GSH) and increasing LIP [29]. Together, these findings indicate that targeting NRF2 promotes ferroptosis by modulating iron metabolism. In the present study, we demonstrate that CD44hi cells, characterized by intrinsically high intracellular iron levels, are particularly susceptible to NRF2 inhibition-induced ferroptosis. Given that NRF2 directly regulates the transcription of SLC7A11 and GPX4, two essential components of the X_c_ ^−^/GSH/GPX4 antioxidant axis [30], inhibition of NRF2 may suppress cystine import and glutathione synthesis, leading to impaired detoxification of lipid peroxides. This loss of redox control, together with NRF2-dependent dysregulation of iron metabolism and expansion of the labile iron pool, is likely to cooperatively enhance lipid peroxidation and drive ferroptotic cell death in CD44hi cells.

There are several limitations to this study. In the LAPC9 model, we were unable to clearly resolve discrete CD44+ and CD44- subpopulations. To address this, we employed negative staining as a gating strategy and sorted the top 30% CD44hi and bottom 30% CD44low cells. Although *in vitro* assays and RNA-seq analyses revealed distinct transcriptional and functional features between these subpopulations, the exclusion of intermediate CD44-expressing cells may have affected our overall findings. Future single-cell RNA-seq with multi-omic analysis of the LAPC9 model will be necessary to more comprehensively characterize CD44-associated heterogeneity and capture the full spectrum of cellular states.

## Material and Methods

### Bioinformatic analysis

Single-cell RNA-seq data was from GEO database GSE210358 CRPC patient1 (GSM6428952_1778) and GSE137829 CRPC patient5 (GSM4711414_P5). The were processed using Seurat. Cells were filtered by gene counts and mitochondrial content, normalized, and integrated using canonical Seurat workflows. Cells were filtered to remove low-quality cells and doublets by retaining cells with 200-6,000 detected genes and <10% mitochondrial gene expression. The remaining cells were normalized using LogNormalize, and datasets were integrated using canonical Seurat integration workflows. Dimensionality reduction (PCA, UMAP) and clustering (SNN graph) were performed with default parameters. Cluster markers were identified using the Wilcoxon test with FDR correction, and cell types were assigned based on canonical markers. Cell types were annotated using the SingleR package, employing a reference-based classification approach. Cells annotated as epithelial were selected and retained for downstream analyses, while non-epithelial cell populations were excluded.

Bulk RNA-seq data were trimmed and aligned to the human reference genome (GRCh38) using STAR, and gene-level read counts were quantified using featureCounts. Differential expression analysis was performed in R (v4.5.1) using DESeq2 (v1.48.1). Genes with an adjusted P value < 0.05 and |log₂ fold change| ≥ 1 were considered significantly differentially expressed. Log₂ fold change shrinkage was performed using apeglm (v1.30.0).

Public bulk RNA-seq datasets were downloaded from the GEO database and processed either from raw FASTQ files using the same alignment and quantification pipeline or from provided count matrices using consistent normalization and statistical modeling. Batch effects arising from the integration of multiple datasets were corrected using ComBat implemented in the sva package within the Bioconductor framework.

Differentially expressed genes were subjected to pathway enrichment analysis using clusterProfiler, including Gene Ontology (GO), KEGG, Reactome, and MSigDB Hallmark pathways. Gene set enrichment analysis (GSEA) was performed using clusterProfiler. Master regulator analysis was conducted using VIPER to infer transcription factor activity based on context-specific regulatory networks. Data visualization was performed using ggplot2 (v4.0.1) and EnhancedVolcano (v1.26.0). Bulk RNA-sequencing data and corresponding clinical annotations for prostate cancer patients were obtained from The Cancer Genome Atlas (TCGA-PRAD) and the DKFZ prostate cancer cohort. Gene expression data were normalized as transcripts per million (TPM) and log₂-transformed for downstream analyses. Spearman rank correlation analysis was performed to evaluate the relationship between the SCL score and iron score, as well as the expression levels of KDM3A and NFE2L2 (NRF2).

To determine the prognostic significance of CD44-associated iron metabolic features, a FerroScore was constructed using Cox proportional hazards regression. Expression levels of CD44, FTL, FTH1, SLC11A2, KDM3A, and NFE2L2 were included as covariates in a univariate Cox regression model for biochemical recurrence or progression-free survival. Regression coefficients (β) derived from the Cox model were used to calculate a risk score for each patient. Patients were stratified into high- and low-FerroScore groups using the median FerroScore as the cutoff. Biochemical recurrence–free survival and progression-free survival were analyzed using Kaplan–Meier methods, and statistical significance between groups was assessed using the log-rank test.

Gene signatures including: Iron metabolism, androgen receptor (AR), Wnt, neuroendocrine (NE), stem cell-like (SCL), and NRF2 activity score were showed in supplementary figure 1. Signature scores were compared between subpopulations using a two-tailed unpaired Student’s t-test. Statistical significance was visualized on box-and-jitter plots generated with the ggplot2 package in R, with significance levels represented by asterisks according to the following thresholds: P<0.001, P<0.01*, P<0.05, and ns (not significant). All comparisons were conducted using the stat_compare_means function from the ggpubr package.

### Cell line culture

Human osteotropic PCa PC-3M-Pro4-luc2 [16] and 22RV1 cells [17] were maintained as described previously [18]. PC-3M-Pro4-luc2, PC3-shCD44 and PC3-shCD44-GFP cells cultured in media (Dulbecco’s modified Eagle’s medium with 10% fetal calf serum II and 1% penicillin–streptomycin). 22RV1, 22RV1-CD44hi, 22RV1-shCD44 and 22RV1-shCD44-GFP cultured in media (RPMI 1640 with 10% fetal calf serum, 1% penicillin–streptomycin and 1% L-glutamate). Cells were maintained at 37 °C with 5% CO2.

### Reagent conditions

BODIPY 581/591 C11 (C11-BODIPY, ThermoFisher Scientific, D3861, 10uM, 30min), Brusatol (Sigma-Aldrich, SML1868), Deferoxamine (DFO, Sigma-Aldrich, D9533, 10uM, 72h), FerrOrange (Dojindo, NC1808697, 1uM, 30min), Ferrostatin-1 (Sigma, SML0583, 10uM, 24h), Iron(III) nitrate (Fe(NO3)3·9H2O (5uM, 72h), Liproxstatin-1 (Sigma, SML1414-5MG, 5 µM, 24h). CBA-1 (ProbChem, PC-20353, 2uM, 72h).

### Cell viability

Cell viability was evaluated by plating 40,000 cells/well in 96-well plates using PrestoBlue viability assay according to the manufacturer’s protocol. Cells were treated with different concentrations of Brusatol as indicated for 72 h. PrestoBlue reagent (ThermoFisher, P50201) was added after 72 h treatment and cells were incubated for 1 h before recording fluorescence intensities (ex. 560 nm; em. 590 nm) using a Varioskan Lux Microplate Reader.

### Flow Cytometry and cell sorting

Single cells obtained from digested tissues or cell lines were washed using FACS buffer (0.5% BSA, 2 mM EDTA in PBS, pH 7.4). The cells were then resuspended in 100 µl of FACS buffer containing anti-CD44-APC (1:20 dilution, BD Biosciences, clone G515) and incubated for 30 minutes in the dark at room temperature. Following incubation, the cells were washed again with FACS buffer and resuspended in 500 µl of FACS buffer per tube. The suspension was supplemented with DAPI at a final concentration of 5 µg/mL and kept on ice until analysis. FACS analysis and cell sorting were conducted using a BD LSRII flow cytometer (BD Biosciences).

### Generation of GFP-Luciferase-labelled PDX models

For the generation of GFP-Luciferase labeled LAPC9 PDX models, MACS-purified LAPC9 human (tumor) single cells were seeded in collagen-coated plates (rat collagen 0.3mg/ml, R&D Systems, 3447-020-01) and transduced with GFP-Luc lentiviral vectors (ATG-2460; GFP-click beetle luciferase) at a multiplicity of infection (MOI) of 1. After 2 days, LAPC9-GFP-Luc tumor cells were transferred in 3D culture conditions to allow organoid formation. Once formed, organoids were resuspended in Matrigel (Corning) and subcutaneously injected into mice (under anesthesia). Tumor growth was monitored through the IVIS Imaging System (PerkinElmer) following intraperitoneal injection of D-sodium salt luciferin (150mg/kg). To deplete potential non-transduced tumor cells, the newly generated LAPC9-GFP-Luc tumors were collected at the endpoint, digested, and FACS was performed to select the GFP+ cells. FACS-purified GFP+ cells were then re-injected subcutaneously into new mouse hosts.

### Organoid Culture

Tissues were collected in a basic medium [Advanced D-MEM/F-12 (ThermoFisher Scientific) supplemented with 1 ml Primocin (Invivogen), 1% GlutaMAX, and 10 mM HEPES (ThermoFisher Scientific)]. The tissues were finely minced with a scalpel and incubated at 37°C in a solution containing 5 mg/ml collagenase type II (Gibco), 15 µg/ml DNase I (Sigma-Aldrich), and 10 mM Y-27632, with occasional mixing for 1–3 hours until fully digested. The resulting cell suspension was centrifuged at 700 rcf for 5 minutes and washed with the basic medium. The cell pellet was subsequently incubated at 37°C in 2 ml of TripLE Express (ThermoFisher Scientific) for 15 minutes, with pipetting every 5 minutes to aid digestion. The digested suspension was passed through a 50 µm-pore size strainer (Celltrics, Sysmex) and washed with the basic medium. When necessary, cells were treated with erythrocyte-lysing buffer for 5 minutes to remove red blood cells, followed by another wash with the basic medium. Cell counting was performed using trypan blue and an automated cell counter (TC20, Bio-Rad). Afterward, cells were centrifuged and resuspended in complete prostate cancer organoid medium (prepared as described previously [19]) at a concentration of 500,000 cells/ml. The suspension was seeded at a volume of 1.5 ml per well in 6-well ultra-low attachment plates (ULA plates, Corning). Fresh medium was added every 2–3 days until the organoids were ready for downstream applications.

### Ex vivo Tissue Culture

Freshly harvested LAPC9 tissues were aseptically cut into 1 mm-thick serial slices. These tissue slices were then carefully positioned on the membrane of a 0.4 µM pore polypropylene 6-well transwell (ThinCert, Grainer Bio-one International) and cultured in 1.5 ml of DMEM supplemented with 10% FBS, 1% penicillin/streptomycin, and the specified compound at 37°C for 4 days. Before incubation, the plate containing the tissue slices was placed inside a sealed chamber and flushed with oxygen (3 L/min) for 3 minutes. After 4 days of culture, the tissues were collected and processed for flow cytometry analysis and Western blot.

### Western Blot

Cells were lysed in RIPA buffer (50 mM Tris-HCl pH 8.0, 100 mM NaCl, 5 mM EDTA, 0.2% SDS, 0.5% sodium deoxycholate, 1% Triton-X100) supplemented with protease and phosphatase inhibitors (cOmplete Mini protease inhibitor cocktail and PhosStop, both from Merck Millipore). Tissue samples were homogenized in RIPA buffer using a metallic bead and the TissueLyser II (Qiagen) for 1 cycle of 2 minutes at 20 Hz. Organoids were resuspended in 150 µl of RIPA buffer and homogenized with a 0.3 ml syringe. Homogenized samples were centrifuged at >16,000 g for 15 minutes at 4°C, and the supernatants were collected. Protein concentrations were determined using the Bradford assay, and 10–30 µg of each sample was loaded for SDS-PAGE. Proteins were then transferred onto Immobilon-PVDF membranes (Millipore). Blots were incubated overnight with primary antibodies, followed by washing with PBS-Tween (refer to Supplementary Table 1 for details on the antibodies used). The blots were then incubated with either anti-mouse or anti-rabbit secondary antibodies. After additional PBS-Tween washes, the proteins were visualized using the Supersignal ECL detection kit (ThermoFisher) and imaged with the Amersham Imager 600 (GE).

### Western blot quantifications

Western blot signals were quantified using ImageJ. Equal areas were defined for each lane and quantified using the Plot Lanes function in ImageJ. Signals arising from individual antibodies were normalized against the antibody signal of the loading control.

### RNA isolation and real-time quantitative PCR

Total RNA was extracted using Trizol Reagent (Invitrogen), and cDNA was synthesized via reverse transcription following the manufacturer’s protocol (Promega). For qPCR analysis, 10 ng of cDNA per reaction was amplified using the CFX Real-Time Detection System (Bio-Rad, Cressier, Switzerland) with SYBR Green Supermix reagent (BioRad). Gene expression levels were normalized to the reference transcripts HPRT and ACTB. The primer sequences used for the reactions are provided in Supplementary Table 2.

### Suppressing CD44 with shRNAs

shRNA for CD44 (TRCN0000296191, TRCN0000289233 and TRCN0000308110) were obtained from Sigma’s MISSION library and shRNA for CD44 with eGFP was from Addgene (Plasmid#19123). Lentiviral infection and selection were performed as described previously [16].

### Bulk RNA barcoding and sequencing (BRB-seq)

Bulk transcriptomic analysis was conducted on sorted LAPC9 CD44hi and CD44low cells. RNA was extracted using Alithea Isolation Reagent. Only RNA samples with quantities between 200 ng and 1000 ng, and an RNA Integrity Number (RIN) greater than 6, were included. The samples were then sent to Alithea Genomics SA (Lausanne, Switzerland) for library preparation and sequencing. For prostate tissue samples, highly multiplexed 3’-end bulk RNA barcoding and sequencing (MERCURIUS™ BRB-seq) was used, while for organoids, extraction-free MERCURIUS™ DRUG-seq was utilized. Library preparation followed the manufacturer’s protocol for the MERCURIUS™ BRB-seq (prostate tissue) or DRUG-seq (organoids) kits for Illumina sequencing. Sequencing was performed on the Illumina NovaSeq 6000 platform. RNA-seq raw counts were analyzed using DESeq2, with genes retained if they had at least 3 UMI counts in at least 3 samples. Differential gene expression was determined using an adjusted p-value threshold of 0.1. Functional enrichment analysis was performed using Enrichr.

### Immunohistochemistry (IHC) Stainings

Sections, 4 µm thick, were cut from FFPE blocks, stained with hematoxylin and eosin (H&E), and mounted using Entellan (Merck Millipore). For CD44 staining, the sections underwent antigen retrieval by heating in a pressure cooker for 10 minutes in citrate buffer (pH 6.0). After cooling, sections were thoroughly washed under running water. To block endogenous peroxidases, sections were incubated with 3% H2O2 at room temperature for 15 minutes, followed by two washes with PBS. Blocking was then performed using a 3% BSA solution in PBS-Tween 20 (0.1%, PBS-T) for 1 hour at room temperature. The sections were subsequently incubated overnight with 100 µl of anti-CD44 antibody (1:100, mouse) or mouse IgG as a control, as listed in Supplementary Table 3. After overnight incubation, the sections were washed once with PBS-T and twice with PBS. They were then incubated with 100 µl of EnVision anti-mouse reagent (Agilent Technologies) for 30 minutes. Following this, sections were again washed once with PBS-T and twice with PBS before development in freshly prepared AEC solution (Dako) until the desired signal was achieved. The sections were washed with water, counterstained with hematoxylin, and mounted with Entellan. Finally, slides were digitized using the Panoramic 250 Flash III slide scanner (3D Histech).

### Immunofluorescence (IF) Staining

FFPE sections (4 μm) were deparaffinized and rehydrated using xylene and ethanol. Antigen retrieval was performed with either a citrate-based buffer (pH 6) (Adipogen, H-3300) or a 10 mM Tris (pH 9)/1 mM EDTA/0.05% Tween buffer. The slides were blocked in a blocking solution (10% donkey serum, 0.1% PBS-Tween or 1% BSA, 0.1% PBS-Tween) for 1 hour at room temperature (RT), followed by overnight incubation at 4°C with primary antibodies (Supplementary Table 3). Secondary antibodies (anti-rabbit/mouse/goat) conjugated to Alexa Fluor®-647, -555, -488 fluorochromes (Life Technologies) were applied for 90 minutes at a 1:250 dilution in the blocking solution. Sections were then counterstained with DAPI (1 μg/ml, ThermoFisher, 62248) for 10 minutes, washed, and mounted with Prolonged Diamond Antifade reagent (ThermoFisher, P36970). Immunofluorescence images were captured using a CellVoyager CQ1 microscope (Yokogawa) and a slide scanner (3DHistech Panoramic 250 Flash II).

### Iron measurement

Intracellular chelatable iron was determined using the fluorescent probe FerroOrange from Dojindo (F374). FerroOrange is a fluorescent probe that enables live-cell fluorescence imaging of intracellular Fe2+. Cells were incubated with 1 μM for 30 min, and data were captured using FACS at PE laser.

### Lipid peroxidation assay

C11-BODIPY581/591 dye (Thermo Fisher Scientific, D3861) was used to detect lipid peroxidation in live cells. For flow cytometry, 500,000 cells were seeded in 6-well plates and allowed to adhere overnight at 37°C. Then, cells were treated with dimethyl sulfoxide (DMSO) or Brusatol (0.1uM) for 24 hours. Cells were then trypsinized, washed, suspended in Hanks’ balanced salt solution (HBSS) containing 2uM C11-BODIPY581/591, and incubated at 37°C for 15 min. Cells were pelleted and resuspended in HBSS. Oxidation of the polyunsaturated butadienyl portion of the dye results in a shift of the fluorescence emission peak from 590 nm to 510nm.

### Chromatin Immunoprecipitation (ChIP) and qPCR

Chromatin immunoprecipitation (ChIP) was performed using a modified protocol based on pervious study. Briefly, cells were crosslinked with 1% formaldehyde for 8 minutes at room temperature, followed by quenching with 125 mM glycine for 5 minutes.

Samples were washed with cold PBS and lysed to obtain nuclei. Chromatin was fragmented by sonication to an average size of 200 – 500 bp. For each immunoprecipitation, equal amounts of chromatin were incubated overnight at 4°C with H3K9me2 antibody [Abcam, ab1791, 1:100] or normal Rbt IgG as a negative control. Antibody-chromatin complexes were captured using protein A/G magnetic beads, washed sequentially with low-salt, high-salt, LiCl, and TE buffers, and eluted according to kit instructions. Crosslinks were reversed at 65°C for 4 hours, followed by treatment with RNase A and proteinase K. DNA was purified using spin columns.

Purified ChIP DNA was analyzed by quantitative PCR (qPCR) using SYBR Green Master Mix (BioRad). Primers were designed to amplify target promoter or enhancer regions of interest, with input DNA used for normalization. Data were expressed as percent input or fold enrichment over IgG using the ΔCt method.

### Animals Maintenance and in vivo Experiment

Animal experiments were carried out in accordance with Bern cantonal guidelines. Mice had unrestricted access to food and fresh water and were housed with a maximum of 5 animals per cage. For xenograft surgery, eighteen 5-week-old male CB17/SCID mice were anesthetized via subcutaneous injection with a cocktail of medetomidine (Dorbene) 1 mg/kg, midazolam (Dormicum) 10 mg/kg, and fentanyl 0.1 mg/kg. Under sterile conditions, the sorted LAPC9 cells were grouped into 2 categories: GFP+ CD44hi, GFP+ CD44low. A total of 100,000 single cells were suspended in 50 µl of Matrigel and injected subcutaneously into the mice. Anesthesia was reversed by subcutaneous injection of atipamezol (Revertor R) 2.5 mg/kg and flumazenil (Anexate R) 0.5 mg/kg, along with buprenorphine (Temgesic) 0.1 mg/kg for analgesia. The sutured wound was disinfected using an iodopovidone solution. Mice were monitored twice a week for changes in body weight, tumor size, and any signs of acute toxicity. Tumor size was assessed by palpation and compared to standardized size beads to minimize discomfort during the experiment. Mice were euthanized as soon as signs of acute toxicity were observed or when the tumor size reached 8 mm.

The tumor growth was monitored by using an in vivo imaging system equipped with integrated micro-CT (IVIS-CT). Mice were anesthetized with 3% isoflurane in oxygen and maintained under anesthesia throughout image acquisition. Prior to imaging, animals received an intraperitoneal injection of D-luciferin potassium salt (150 mg/kg) dissolved in sterile PBS. Imaging was initiated 15 minutes post-injection to capture peak bioluminescent signal. Images were analyzed using Living Image software. Regions of interest (ROIs) were manually drawn over the site of luciferase expression, and corresponding signals were quantified as total flux (photons/sec) after background subtraction. CT images were used for anatomical reference and to ensure accurate ROI placement.

Ten CB17/SCID mice were subcutaneously implanted with LAPC9-GFP tumor pieces and randomly assigned into two groups: vehicle control (DMSO) and Brusatol treatment. Treatment began 1 week after tumor implantation. Mice in the Brusatol group received Brusatol at 2 mg/kg via intraperitoneal injection every other day for 3 weeks, while control mice received an equivalent volume of DMSO. Tumor progression was monitored using an IVIS-CT imaging system, and bioluminescent signal intensity was used to assess tumor growth longitudinally. When tumors reached 8 mm in diameter, all mice were euthanized and tumors were excised. Tumor size and tumor weight were measured immediately after collection.

### Statistic

Statistical analyses were conducted using Kaplan-Meier (K-M) analysis, unpaired t-test, chi-square test, one-way ANOVA, and two-way ANOVA with multiple comparison testing, as specified in each figure legend. All statistical analyses and data visualizations were performed using GraphPad Prism 10 software (GraphPad Software Inc.) or RStudio version 2023.06.1, unless otherwise noted. Data are expressed as mean ± SD from at least three independent experiments, unless indicated in the figure legend. Statistical significance is indicated as follows: * p ≤ 0.05, ** p ≤ 0.01, *** p ≤ 0.001, and **** p ≤ 0.0001.

## Reference

1. Easwaran H, Tsai H-C, Baylin Stephen B. Cancer epigenetics: tumor heterogeneity, plasticity of stem-like states, and drug resistance. Mol. Cell. (2014).

2. Perlmutter, M. A. & Lepor, H. Androgen deprivation therapy in the treatment of advanced prostate cancer. Rev Urol 9 Suppl 1, S3–8 (2007).

3. Scher, H. I. et al. Trial Design and Objectives for Castration-Resistant Prostate Cancer: Updated Recommendations From the Prostate Cancer Clinical Trials Working Group 3. J Clin Oncol 34, 1402–1418 (2016).

4. Tang F, Xu D, Wang S, Wong CK. Chromatin profiles classify castration-resistant prostate cancers suggesting therapeutic targets. Science. 2022 May.

5. Naor D, Nedvetzki S, Golan I, Melnik L, Faitelson Y. CD44 in cancer. Crit Rev Clin Lab Sci. 2002;39(6):527–79.

6. Chen C, Zhao S, Karnad A, Freeman JW. The biology and role of CD44 in cancer progression: therapeutic implications. J Hematol Oncol. 2018 May 10;11(1):64.

7. Müller, S., Sindikubwabo, F., Cañeque, T. et al. CD44 regulates epigenetic plasticity by mediating iron endocytosis. Nat. Chem. 12, 929–938 (2020).

8. Abbate, V. & Hider, R. Iron in biology. Metallomics Integr. Biometal Sci. 9, 1467–1469 (2017).

9. Farida B, Ibrahim KG, Abubakar B, Malami I, Bello MB, Abubakar MB, Abbas AY, Imam MU. Iron deficiency and its epigenetic effects on iron homeostasis. J Trace Elem Med Biol. 2023 Jul;78:127203. doi: 10.1016/j.jtemb.2023.127203.

10. Ghoochani A, Hsu EC, Aslan M, Rice MA, Nguyen HM, Brooks JD, et al. Ferroptosis inducers are a novel therapeutic approach for advanced prostate cancer. Cancer Res. 2021;81:1583–94.

11. Wang, Y., Ma, Y. & Jiang, K. The role of ferroptosis in prostate cancer: a novel therapeutic strategy. Prostate Cancer Prostatic Dis 26, 25–29 (2023).

12. Joseph M. Chan et al., Lineage plasticity in prostate cancer depends on JAK/STAT inflammatory signaling. Science377,1180–1191(2022).

13. Dong, B., Miao, J., Wang, Y. et al. Single-cell analysis supports a luminal-neuroendocrine transdifferentiation in human prostate cancer. Commun Biol 3, 778 (2020).

14. Karkampouna, S., La Manna, F., Benjak, A. et al. Patient-derived xenografts and organoids model therapy response in prostate cancer. Nat Commun 12, 1117 (2021).

15. Padeken, J., Methot, S.P. & Gasser, S.M. Establishment of H3K9-methylated heterochromatin and its functions in tissue differentiation and maintenance. Nat Rev Mol Cell Biol 23, 623–640 (2022).

16. Li, Q., Qin, K., Tian, Y. et al. Inhibition of demethylase by IOX1 modulates chromatin accessibility to enhance NSCLC radiation sensitivity through attenuated PIF1. Cell Death Dis 14, 817 (2023).

17. Fan, L., Xu, S., Zhang, F. et al. Histone demethylase JMJD1A promotes expression of DNA repair factors and radio-resistance of prostate cancer cells. Cell Death Dis 11, 214 (2020).

18. Wen Zhang, Vitaliy M. Sviripa, Epigenetic Regulation of Wnt Signaling by Carboxamide-Substituted Benzhydryl Amines that Function as Histone Demethylase Inhibitors, iScience, Volume 23, Issue 12, 2020, 101795, ISSN 2589-0042.

19. Zhang, D.D. Thirty years of NRF2: advances and therapeutic challenges. Nat Rev Drug Discov 24, 421–444 (2025).

20. Cai, S.J., Liu, Y., Han, S. et al. Brusatol, an NRF2 inhibitor for future cancer therapeutic. Cell Biosci 9, 45 (2019).

21. Annadurai Anandhan et al.,NRF2 controls iron homeostasis and ferroptosis through HERC2 and VAMP8.Sci. Adv.9,eade9585(2023).

22. Zhang, S., Xin, W., Anderson, G.J. et al. Double-edge sword roles of iron in driving energy production versus instigating ferroptosis. Cell Death Dis 13, 40 (2022).

23. Patrawala L, Calhoun-Davis T, Schneider-Broussard R, Tang DG. Hierarchical organization of prostate cancer cells in xenograft tumors: the CD44+alpha2beta1+ cell population is enriched in tumor-initiating cells. Cancer Res. 2007 Jul 15;67(14):6796–805. Erratum in: Cancer Res. 2007 Sep 15;67(18):8973. PMID: 17638891.

24. Solier, S., Müller, S., Cañeque, T. et al. A druggable copper-signalling pathway that drives inflammation. Nature 617, 386–394 (2023).

25. Wilson S, Fan L, Sahgal N, Qi J, Filipp FV. The histone demethylase KDM3A regulates the transcriptional program of the androgen receptor in prostate cancer cells. Oncotarget. 2017 May 2;8(18):30328–30343.

26. Hansen AM, Ge Y, Schuster MB, Pundhir S, Jakobsen JS, Kalvisa A, Tapia MC, Gordon S, Ambri F, Bagger FO, Pandey D, Helin K, Porse BT. H3K9 dimethylation safeguards cancer cells against activation of the interferon pathway. Sci Adv. 2022 Mar 18;8(11):eabf8627.

27. Ryoo IG, Choi BH, Ku SK, Kwak MK. High CD44 expression mediates p62-associated NFE2L2/NRF2 activation in breast cancer stem cell-like cells: Implications for cancer stem cell resistance. Redox Biol. 2018 Jul;17:246–258.

28. Lim, P.J., Duarte, T.L., Arezes, J. et al. Nrf2 controls iron homoeostasis in haemochromatosis and thalassaemia via Bmp6 and hepcidin. Nat Metab 1, 519–531 (2019).

29. Zhu X, Huang N, Ji Y, Sheng X, Huo J, Zhu Y, Huang M, He W, Ma J. Brusatol induces ferroptosis in oesophageal squamous cell carcinoma by repressing GSH synthesis and increasing the labile iron pool via inhibition of the NRF2 pathway. Biomed Pharmacother. 2023 Nov;167:115567.

30. Jiang, X., Yu, M., Wang, Wk., et al. The regulation and function of Nrf2 signaling in ferroptosis-activated cancer therapy. Acta Pharmacol Sin 45, 2229–2240 (2024).

