## Supplemental Figure for "Iron Metabolism as a Therapeutic Vulnerability in Stem Cell-Like Castration-Resistant Prostate Cancer"

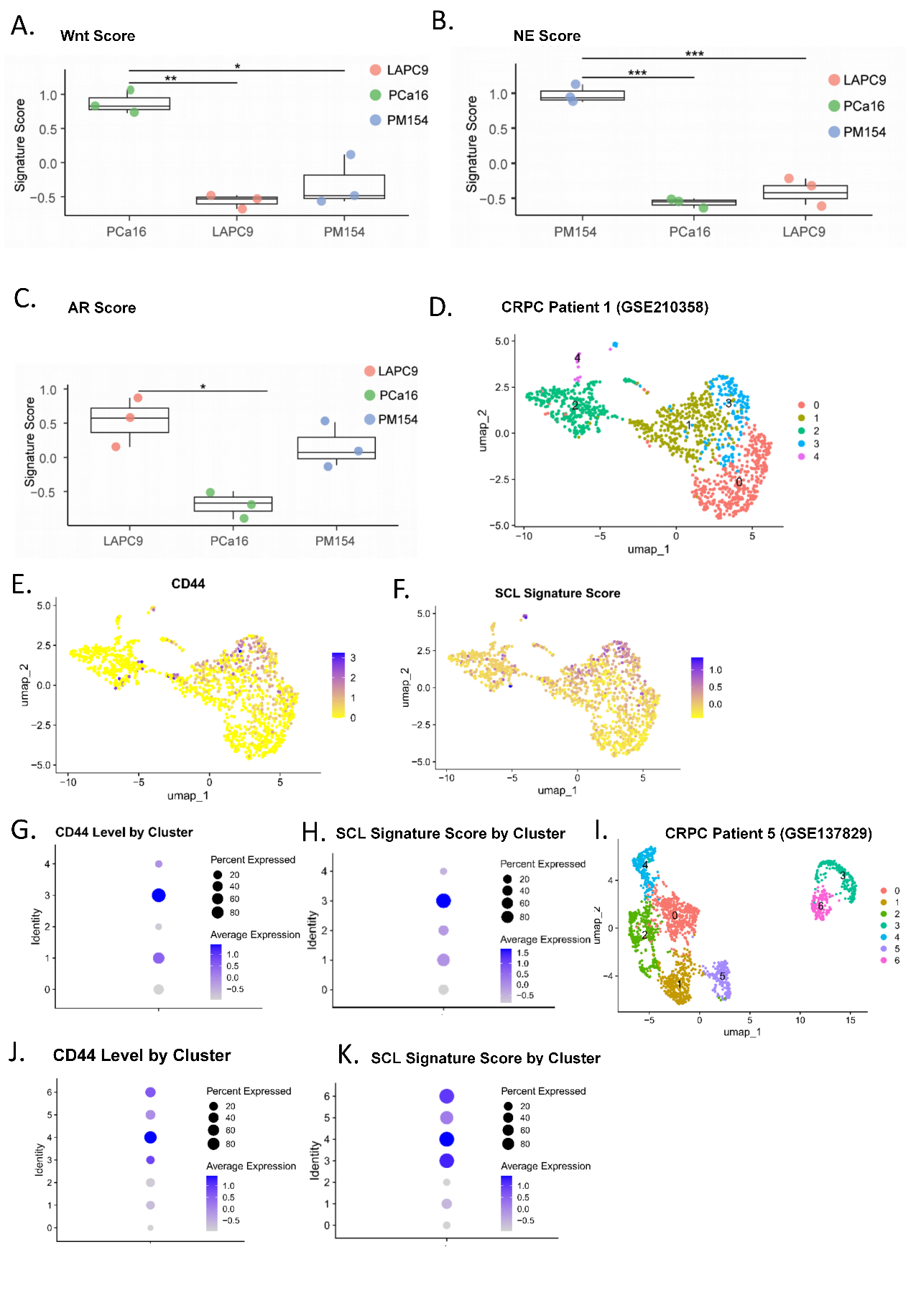

**Supplement Figure 1. A-C.** Signature analysis was identified NE score (A), Wnt score (B) and AR score (C) base on RNA-seq data among LAPC9, PCa16 and PM154 organoids. **D.** Dimensionality reduction of single-cell distribution of CRPC patient 1 (GSE210358) using UMAPs and subsequent identification of cell clusters performed using Seurat workflow. Each cluster of cancer cells were shown in UMAP. **E-H**. CD44 distribution in UMAP shown in (**E**), CD44 level in each cluster shown in (**G**). Signature analysis was identified stem cell like (SCL) score, the SCL score distribution in UMAP shown in (**F**), SCL level in each cluster shown in (**H**). **I.** Dimensionality reduction of single-cell distribution of CRPC patient 5 (GSE137829) using UMAPs and subsequent identification of cell clusters performed using Seurat workflow. Each cluster of cancer cells were shown in UMAP. **J**. CD44 level in each cluster shown by dot plot. **K.** Signature analysis was identified stem cell like (SCL) score, SCL level in each cluster shown by dot plot.

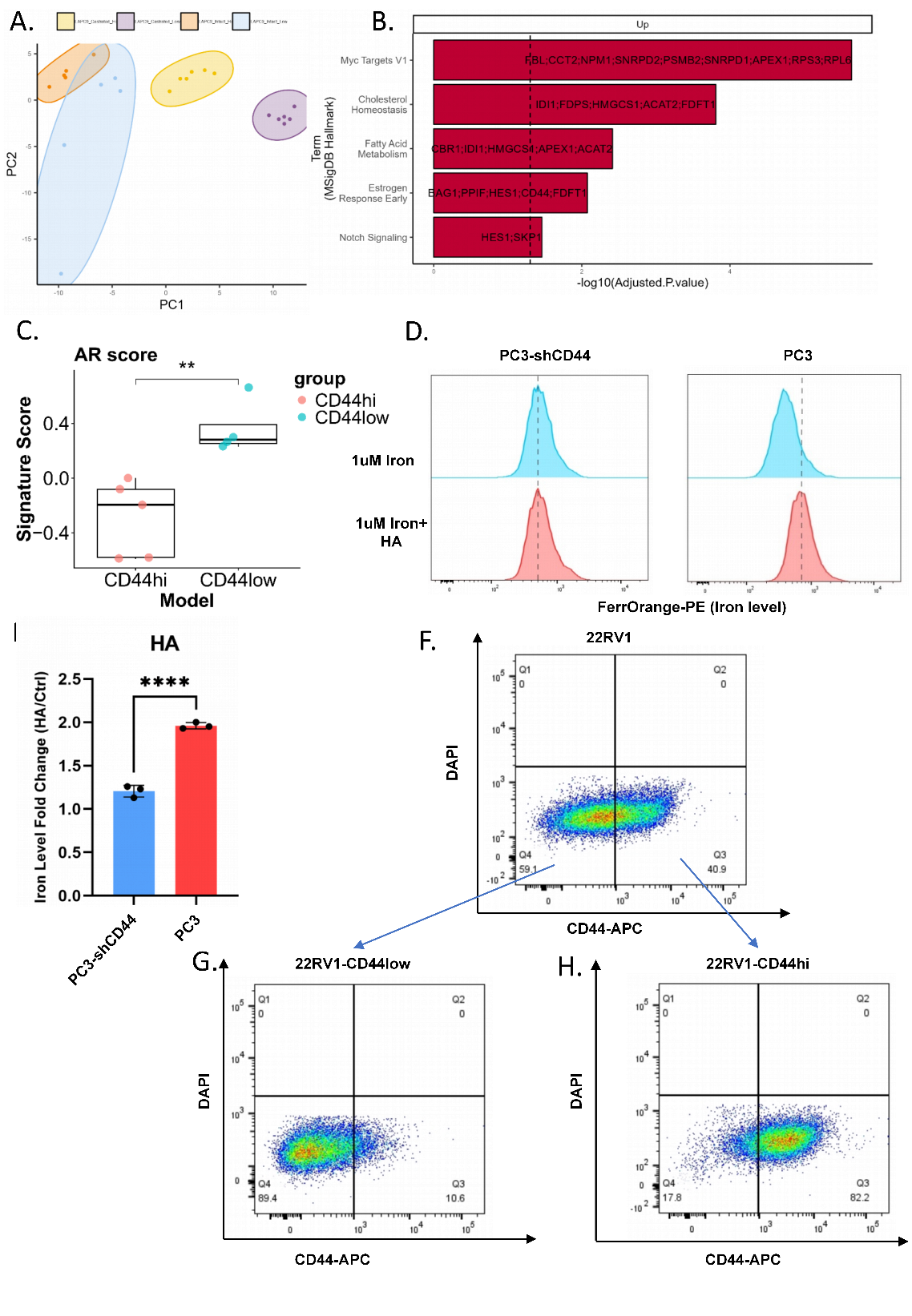

**Supplement Figure 2. A.** RNA-seq performed on CD44hi and CD44low cells from LAPC9 models under intact and castration conditions. Principal component analysis (PCA) plot show the transcriptional difference between CD44hi and CD44low subpopulations. **B.** Pathway enrichment analysis was identified significant pathways in LAPC9 CD44hi subpopulation. **C.** Signature analysis was identified AR score between CD44hi and CD44low cells. **D-E.** Intercellular iron level between two models. CD44hi model PC3 and PC3-shCD44 were treat with iron (1uM) with or without HA (1mg/ml) for 48h. FACS performed on PC3 and PC3-shCD44 cells staining with FerrOrange. **F-H.** FACS sorting CD44hi cell and CD44low cells from 22RV1 cells (**F**). Kept culture as cell line, named 22RV1-CD44low (**G**) and 22RV1-CD44hi (**H**) cell line.

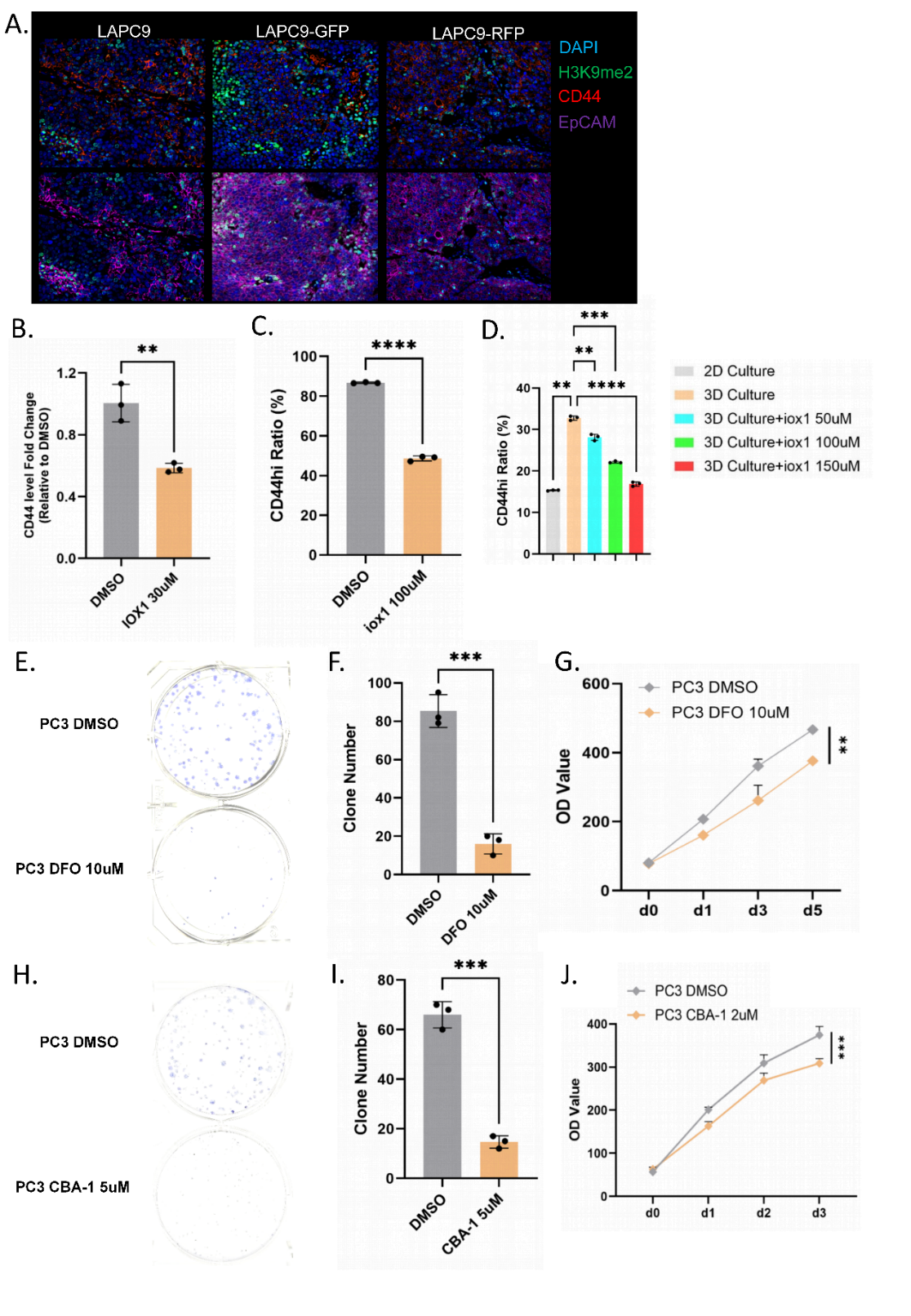

**Supplement Figure 3. A.** Immunofluorescence (IF) staining were performed on LAPC9, LAPC9-GFP and LAPC9-RFP tumors to check CD44 and H3K9me2. **B.** qPCR to compare CD44 between 22RV1-CD44hi cells and treated with 30uM IOX1 for 72h. **C.** FACS staining with CD44-APC was performed on 22RV1 cells treated with or without IOX1 (100uM) for 72h. Ratio of CD44hi cells were compared between two groups. **D.** FACS for 22RV1-CD44low cells cultured in 2D, 3D organoid medium condition and with treat CBA-1 for 3 days. **E.** Clonogenic assay of PC3 cells treated with DMSO or DFO 10uM for 72 h before seeding (500 cells/well). **F.** Measured and quantified the clone number by ImageJ. **G.** Measurement growth curve of PC3 treated with DMSO or DFO 10uM, viability measured by PrestoBlue (n=4). **H.** Clonogenic assay of PC3 cells treated with DMSO or CBSA-1 5uM for 72 h before seeding (500 cells/well). **I.** Meaure and quantified the clone number by ImageJ. **J.** Measurement growth curve of PC3 treated with DMSO or CBA-1 2uM, viability measured by PrestoBlue (n=5).

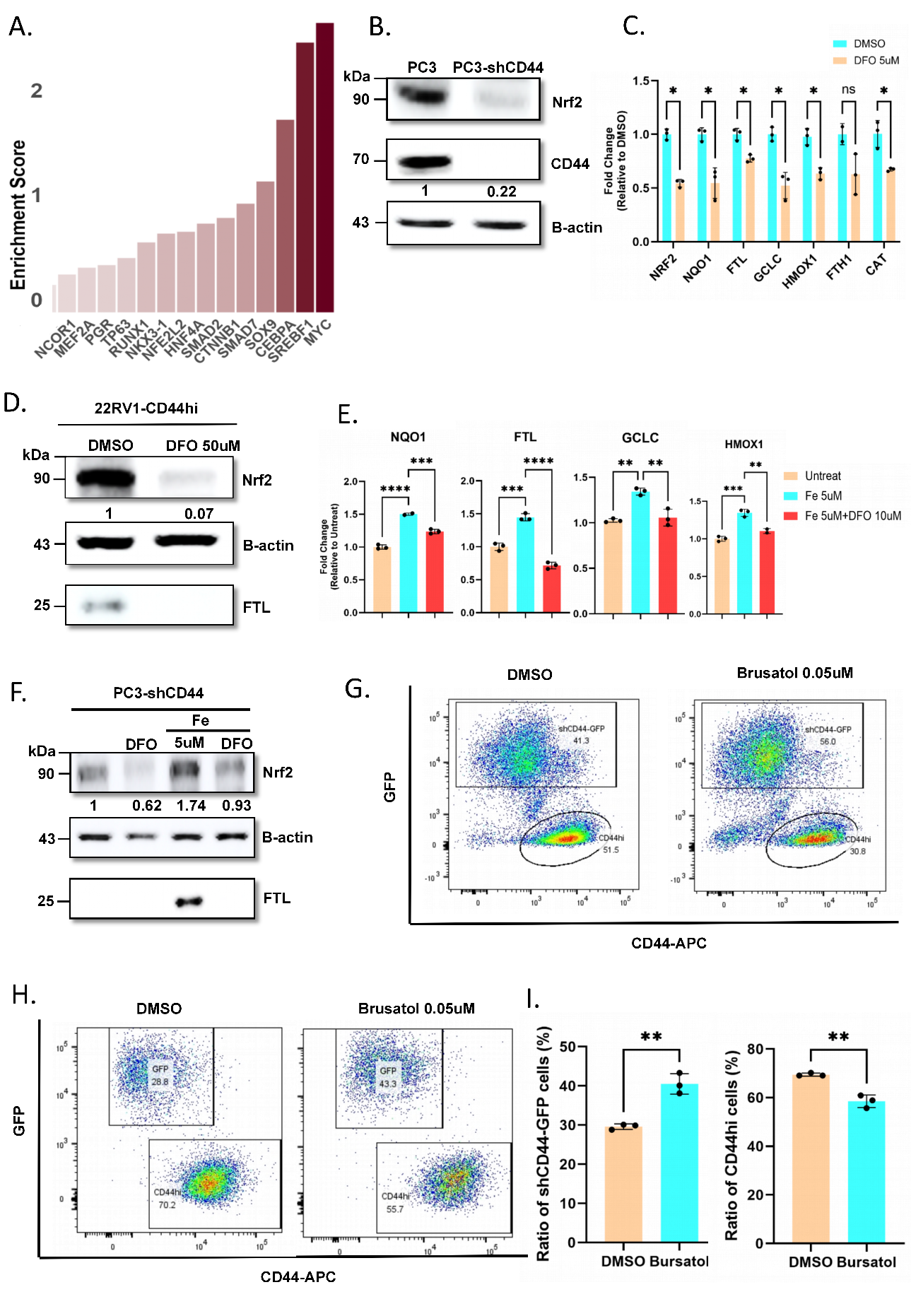

**Supplement Figure 4. A.** RNA-seq performed on CD44hi and CD44low cells from LAPC9 models under intact conditions. Differential expression gene (DEGs) analysis between sorted CD44hi cell and CD44low cells of LAPC9 PDX. Identified master regulators, which could regulate DEGs, in CD44hi subpopulation versus the CD44low was performed by VIPER analysis. **B.** Western-blot confirming CD44 and NRF2 level in 22RV1-CD44hi treat with or without DFO 50uM for 72h. The b-Actin was used as loading control. Signal of protein were quantified by Image J. **C.** qPCR for checking mRNA level of NRF2 downstream genes between 22RV1-CD44hi treat with or without DFO 5uM (3 replicates per sample). **D.** Western Blotting analysis of NRF2 and FTL levels. Protein detection signals were quantified by ImageJ. PC3 cells were treated with iron (2uM) and iron chelator DFO (10uM) for 72h. **E.** qPCR for mRNA level of NRF2 downstream genes. PC3 cells treated with Iron (5uM)+DFO (10uM). **F.** Western Blotting analysis for NRF2 protein levels at PC3 cells and PC3-shCD44 cells. **G.** Mix 22RV1-CD44hi cells and 22RV1-shCD44-GFP cells as 1:1 ratio (400,000 cells total/well), treated with 0.05uM Brusatol for 72h. Preformed FACS check ratio of CD44hi and CD44low cells with DMSO group. **H.** Mix PC3 cells and PC3-shCD44-GFP cells as 1:1 ratio (100,000 cells total/well), treated with 0.05uM Brusatol for 72h. Preformed FACS check ratio of CD44hi and CD44low cells with DMSO group. **I.** The GFP+ ratio and CD44hi ratio were compared.

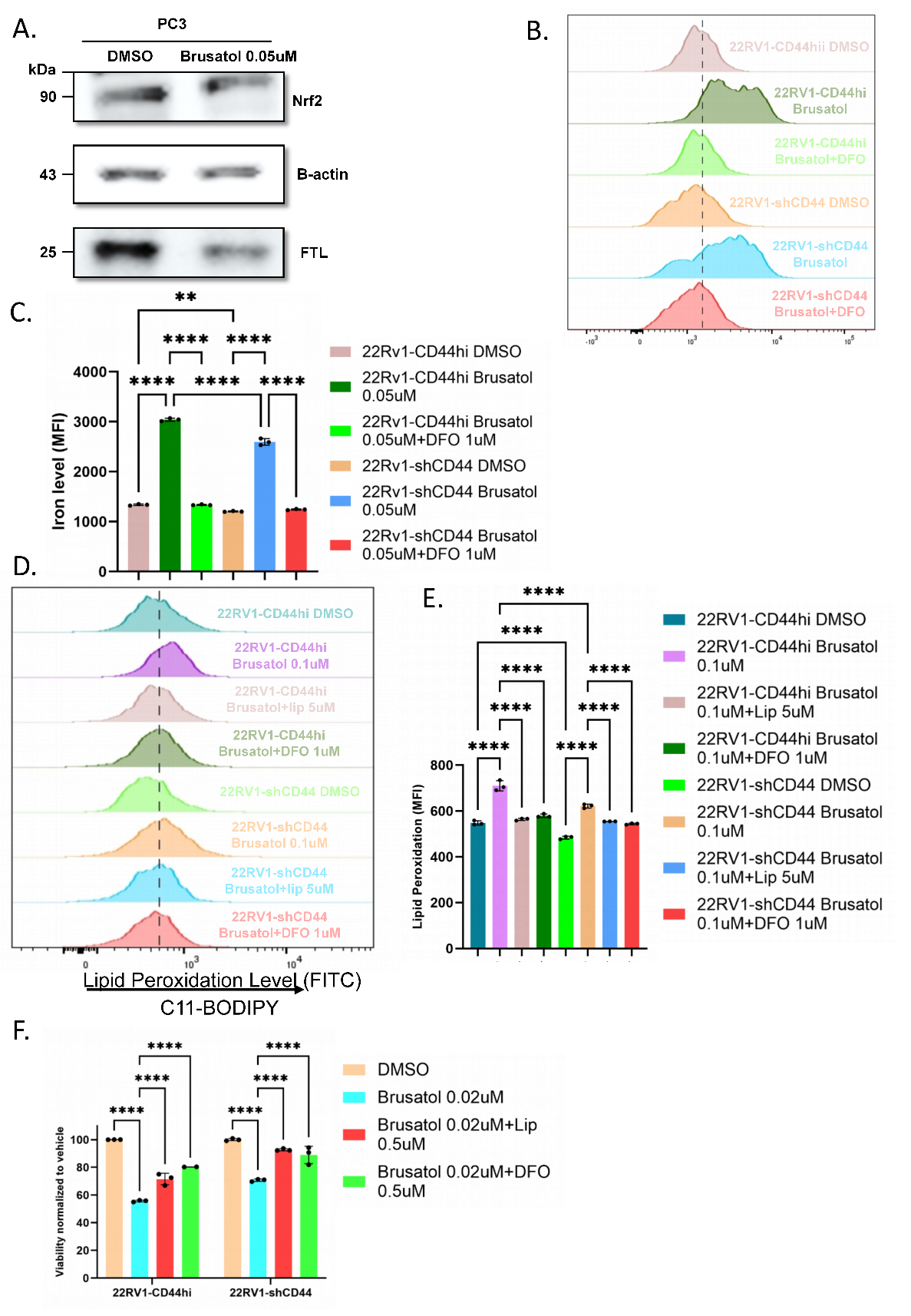

**Supplement Figure 5. A.** Western Blotting analysis for NRF2 and FTL protein levels PC3 cells treated with Brusatol (0.1uM) for 72h. Protein detection signals were quantified by ImageJ. **B, C.** Labile iron pool level after Brusatol treatment. FACS (staining with FerrOrange) on 22Rv1-CD44hi cells and 22Rv1-shCD44 cells treated with Brusatol (0.05uM) with or without DFO (1uM) for 72h (**B**), iron levels were compared among six groups (**C**). **D, E.** Lipid peroxidation levels after Brusatol treatment. FACS (staining with C11-BODIPY) on 22RV1-CD44hi and 22RV1-shCD44 cells treated with Brusatol (0.1uM) with or without Lipostatin-1 (5uM) and DFO (1uM) for 24h (**D**), lipid peroxidation levels were compared among different groups (**E**). **F.** Measurement of 22RV1-CD44hi and 22RV1-shCD44 cells viability by PrestoBlue, Brusatol (0.05uM) treated cells with or without Lipostatin-1 (1uM) and DFO (0.5uM) for 72h (n=3).

**Supplementary Table 1 Gene List for Signature Score**

| Signature Score | Gene List |
| --- | --- |
| Iron Score | "FTL", "FTH", "HMOX1", "TFRC", "SLC11A2", "SLC40A1", "CD44", "NCOA4", "HAMP" |
| SCL Score | "FOSL2", "CD44", "ATF3", "ATXN1", "TACSTD2" |
| AR Score | "AR","DHCR24", "KLK3", "FKBP5", "FOXA1", "NKX3-1" |
| Wnt Score | "TCF7L2", "KLF4", "SOX4", "SNAI1", "TCF7", "KLF2", "AXIN2", "RNF43", "TCF7L2", "CTNNB" |
| NE Score | "NEUROD1", "ASCL1", "TCF12", "MYOG", "MSC", "NHLH1" |
| NRF2 Score | "NQO1", "FTL","FTH1", "CAT", "NFE2L2" |

**Supplementary Table 2 Antibody for Western Blot**

| Antibody | Company | Cat.No | Dilution |
| --- | --- | --- | --- |
| Actin | Cell signaling | 4967 | 1:2000 |
| CD44 Pan specific | BioTechne | BBA10 | 1:1000 |
| c-MYC | Cell signaling | 9402 | 1:1000 |
| Ferritin light chain (FTL) | Proteintech | 10727-1-AP | 1:1000 |
| G9a/EHMT2 Antibody | Cell Signalin | 4658 | 1:1000 |
| H3K9me2 | Abcam | ab1791 | 1:1000 |
| H3 Histon Antibody | Abcam | ab1791 | 1:2000 |
| KDM3A | Proteintech | 12835-1-AP | 1:1000 |
| NRF2 | Proteintech | 16396-1-AP | 1:1000 |
| Anti-Rabbit IgG, HRP-Linked Ab | Cell signaling | 7074 | 1:2000 |
| Donkey anti-mouse IgG (H+L), HRP | ThermoFisher | A16011 | 1:2000 |

**Supplementary Table 3 Primer for qPCR**

| Genes | Forward (5’-3’) | Reverse (5’-3’) |
| --- | --- | --- |
| Actin | GCACAGAGCCTCGCCTT | CCTTGCACATGCCGGAG |
| CAT | AGGGGCCTTTGGCTACTTTG | ACCCGATTCTCCAGCAACAG |
| CD44 | GACAAGTTTTGGTGGCACG | CACGTGGAATACACCTGCAA |
| FTH1 | CATCAACCGCCAGATCAAC | GATGGCTTTCACCTGCTCAT |
| FTL | GCACGGACCCCCATCTCTG | GGCTCAGAAGGCTCTTAGTCG |
| GCLC | TGCCTATGTGGTGTTTGTGGT | ACCACTGCATTGCCACCTTT |
| HMOX1 | GACGGCTTCAAGCTGGTGAT | GCTGGTGTGTAGGGGATGAC |
| HPRT | AGACTTTGCTTTCCTTGGTCAGG | GTCTGGCTTATATCCAACACTTCG |
| NQO1 | TGGAGTCCCTGCCATTCTGA | ACTGCCTTCTTACTCCGGAAGG |
| NRF2 | AGGTTGCCCACATTCCCAAA | ACGTAGCCGAAGAAACCTCATT |

**Supplementary Table 4 Antibody for IHC and IF**

| Antibody | Company | Cat.No | Dilution |
| --- | --- | --- | --- |
| AR | Abcam | ab133273 | 1:100 |
| CD44 | BD Pharmingen | 550988 | 1:100 |
| EpCAM | R&D Systems | AF960 | 1:300 |
| H3K9me2 | Abcam | ab1791 | 1:200 |
| NKX3.1 | AthenaES | 314 | 1:250 |
| IgG Goat | Vector Laboratories |  | 1:1000 |
| IgG Mouse | Vector Laboratories | I-2000-1 | 1:300 |
| IgG Rabbit | Vector Laboratories | I-1000-5 | 1:400 |
| AF488 donkey anti-rabbit | Life technolgoies | A21206 | 1:250 |
| AF555 donkey anti-mouse | Life technolgoies | A31570 | 1:250 |
| AF647 donkey anti-goat | Life technolgoies | A21447 | 1:250 |
